# Marmosets mutually compensate for differences in rhythms when coordinating vigilance

**DOI:** 10.1101/2023.09.28.559895

**Authors:** Nikhil Phaniraj, Rahel K. Brügger, Judith M. Burkart

**Author notes:** These two authors have contributed equally to this work (Nikhil Phaniraj, Rahel K. Brügger).

## Abstract

Synchronisation is widespread in animals, and studies have often emphasised how this seemingly complex phenomenon can emerge from very simple rules. However, the amount of flexibility and control that animals might have over synchronisation properties, such as the strength of coupling, remains underexplored. Here, we studied how pairs of marmoset monkeys coordinated vigilance while feeding. By modelling them as coupled oscillators, we noted that (1) individual marmosets do not show perfect periodicity in vigilance behaviours, (2) even then, pairs of marmosets developed a tendency to take turns being vigilant, a case of anti-phase synchrony, (3) marmosets could couple flexibly; the coupling strength varied with every new joint feeding bout, and (4) marmosets could control the coupling strength; dyads showed increased coupling if they began in a more desynchronised state. Such flexibility and control over synchronisation require more than simple interaction rules. Minimally, animals must estimate the current degree of asynchrony and adjust their behaviour accordingly. Moreover, the fact that each marmoset is inherently non-periodic adds to the cognitive demand. Overall, our study taps into the cognitive aspects of synchronisation and provides a mathematical framework to investigate the phenomenon more widely, where individuals may not display perfectly rhythmic behaviours.

## Introduction

Rhythmic phenomena involving multiple animals are widespread in nature [1–3]. From synchronised flashings of fireflies [4,5], call synchronisation in frogs [6,7] to coupled neuronal activity in bats [8] or synchronised heart rates in humans [9] – synchronicity can be found in a huge variety of organisms in different behavioural contexts, at varying levels and serving multiple functions. Synchronicity not only includes patterns where individuals behave in the same way at the same time, as in the impressive synchronous light flashes of fireflies or coordinated movements of starling flocks [10]. Many phenomena show anti-phase synchronisation in which individuals alternate behaviour [11], as in vocal turn-taking in marmoset monkeys [12,13], meerkats [14], elephants [15] and plain-tailed wrens [16] or gestural exchanges of mother-infant dyads in chimpanzees and bonobos [17]. Whereas in-phase and anti-phase synchrony are the two extremes and frequently occur in nature, synchronisation can, in principle, occur at any phase lag (e.g., [18]).

However, the complexity of these phenomena does not necessarily imply complex driving mechanisms. There is extensive evidence from invertebrates that synchronised patterns can emerge as epiphenomena of animals following very simple rules: local interactions such as flashing sooner than usual when a neighbour flashes in the case of fireflies [19–21]. Such simple mechanistic rules predict patterns of synchronisation that are quite uniform, with minimal variation of rhythmic rates between individuals that are coupled and across behavioural bouts. Animals are also expected to show a limited range of possible adjustments to their inherent rhythm, making it difficult to adjust to more dissimilar stimulus rhythms [3]. This is especially important when looking at contexts such as gait synchronisation [22,23] since the skeletal and motor system inherently limits the movement frequencies of animals [3]. When looking at patterns of behaviour such as gestural exchanges where movements are less constrained, these limiting factors to the flexibility of rhythmic rates might play a lesser role.

What remains underexplored is how flexible the synchronisation patterns are in non-human animals with more complex cognitive abilities. Most studies investigating cognitive mechanisms underlying rhythmic behaviours focus on the perception and production of rhythmic patterns [24]. For example, typical rhythm perception tasks are performed as go/no-go tasks where animals are trained to respond to regular rhythms but not irregular ones and are then confronted with novel regular and irregular sequences (“anisochrony detection”) that they are expected to generalise the patterns of regularity too, i.e., they should respond to regular sequences but not to irregular ones. Both rats [25] and starlings [26,27] are capable of solving such tasks (but not zebra finches: [28] and pigeons: [29]). Other studies use approach behaviour to discern whether animals can discriminate between rhythmic patterns. These tasks seem especially promising when used with mating calls of frogs or insects, where animals seem to show species-specific rhythmic preferences [30,31]. Rhythm production studies in vertebrate animal models rely heavily on training (and rewarding anticipatory movements) and a laboratory setting (i.e., [32–34]. These studies show that primates [34,35], rats [36] and birds [33,37] can successfully achieve synchronisation in tasks where they are required to synchronise a motor action such as pressing a lever or pecking a key to an audio or visual metronome stimulus (even with adaptations to changes in tempo). The most flexible rhythm production has been shown by a sea lion and two parrots who were capable of synchronising to the beat of real music at varying tempos (sea lion: [38,39]; parrots: [40,41]), thus indicating that coupling is not restricted to a certain rate. Even though these lab-based studies offer a very controlled environment to investigate cognitive flexibility of synchronisation abilities, they also heavily rely on motivational factors that do not necessarily relate to how flexible rates of rhythmic abilities can be adjusted. These factors might sometimes be overlooked, especially when encountering negative results [3]. The other avenue that has been taken is relying on more naturalistic observation where rhythmic abilities from a species-specific behavioural repertoire are used (e.g., chimpanzee walking: [42]). Critically, this approach can be further enhanced when combined with mathematical modelling [2].

One such context where animals are expected to show high motivation for temporal coordination, and that is under high selection pressures is anti-predator vigilance. Even without assumptions about organisation at the group level, due to the sheer number of eyes looking out for predators, there is ample evidence that individuals living in bigger groups spend less time being vigilant [43–45]. But in many species, it has been shown that individuals’ vigilance levels are not independent of each other but rather that they are synchronised at the group level (e.g., birds: [46,47]; mammals: [48–50]), leading to periods of higher and lower vigilance. However, vigilance can also be coordinated in anti-phase (i.e., animals maximising the feeding time when any group member is vigilant [51]), mostly achieved via a sentinel system with turn-taking like patterns [52–54], even though evidence for coordination is limited.

For the small, arboreal common marmosets, who are vulnerable to predation in the wild ([55], carnivores: [56], snakes: [57], raptors: [58]), vigilance is a key part of their survival strategy [59]. They follow the general trend of a negative group size effect with bigger groups being associated with lower levels of individual vigilance [60]. Some studies have claimed the presence of a sentinel system [61,62] and more recently marmoset pairs have shown to coordinate their vigilance in a feeding situation, by being more vigilant when the pair mate was feeding than when this was not the case [63]. Furthermore, when feeding in proximity with their mate, they changed their behavioural state more slowly when showing opposite behaviours, i.e., one individual being vigilant and the other feeding [63]. These results suggest temporal coordination between individuals, but how this anti-phase synchrony develops or fades out is currently unknown.

Here, we dynamically modelled the vigilance and feeding behaviours of captive pairs of common marmosets as coupled oscillators using the Kuramoto model. We used a feeding situation where being vigilant and feeding were mutually exclusive (namely, when marmosets were eating mash out of an opaque feeding bowl). First, we described the general properties of marmosets’ vigilance and feeding bouts, especially comparing mean vigilance and feeding durations between times when animals were situated alone versus together on the feeding basket. Next, we modelled the two individuals as coupled oscillators according to the classic Kuramoto model [64,65]. The Kuramoto model is one way to model the development of synchrony and its temporal variations, but it assumes that the individual oscillators are inherently periodic. We thus additionally developed a non-periodic version of the Kuramoto model, as many biological systems are not expected to be inherently periodic [66,67]. Further, we used a control model based on the behaviour (vigilance and feeding) of each of the individuals in the pair when they were situated alone on the feeding basket to model the amount of coupling behaviour that can be expected by chance. By fitting the classic and non-periodic Kuramoto models to the data, we were able to estimate how strongly one individual’s behaviour influenced the other individual, i.e., the coupling strength – through the coupling constant. Positive and negative values of the coupling constant indicate the tendency to reach in-phase and anti-phase synchrony, respectively. We additionally derived the critical coupling constant (the threshold to reach a state of synchrony), using the classic Kuramoto model and the control model. For the classic Kuramoto model, we also obtained the time to reach anti-phase synchrony.

Our predictions are threefold: 1) Most biological oscillators are not perfectly periodic, yet there are abundant examples of synchronisation in the animal world. We do not expect marmoset head oscillations to be periodic in the absence of any input from conspecifics, but we predict that they will still show anti-phase synchrony with a partner, given that the probability of an individual being vigilant when its partner is feeding has been shown to be higher than chance [63,66–68]. 2) If marmosets are following simple, fixed interaction rules to coordinate vigilance, we would expect the coupling strengths to be more-or-less uniform across feeding bouts. Moreover, we would expect the synchronisation dynamics to be uniform. We predict that there will be some variation in the coupling strengths across feeding bouts and that marmosets can synchronise flexibly. 3) If the coupling strength indeed varies across feeding bouts, we predict that the differences in the initial rate of head oscillations can explain some of this variation. This is motivated by the fact that marmosets show sensitivity to initial conditions when synchronising in other modalities. For example, marmosets converge with their pair-mates in acoustic space, i.e., sound more similar to their partner over time, in a process called vocal accommodation [69,70]. During this process, they also move through the acoustic space in a synchronised fashion [71]. The extent of vocal accommodation was proportional to the initial differences in vocal properties, with higher initial differences leading to more vocal accommodation later [72].

## Methods

### Subjects and housing

The behavioural data for this study was collected by Brügger et al. (2023) [63] on five breeding pairs, a pair of adult siblings and a family group with young infants (age of immatures during data collection 1–4 weeks), resulting in a total of 14 adult marmosets (for details on the group composition and testing order see table S1). The animals were housed in heated indoor enclosures with access to outdoor enclosures (during appropriate weather conditions: temperature above 10° C). All enclosures were equipped with climbing materials (natural branches, ropes etc.), resting platforms, hammocks, sleeping boxes, and a bark mulch-covered floor. Feeding of the animals occurred at least twice a day in the mornings (ca. 8:00) with a vitamin enriched mash and during lunch (between 11:00–12:00) with fresh vegetables. In the afternoons, animals received a snack feeding with various additional protein sources (eggs, nuts, insects). Marmosets had access to water ad libitum.

### Procedure

Data was recorded during the regular morning feedings without an experimenter being present to eliminate the effects of vigilance towards the experimenter. Therefore, animals were filmed with one or two video cameras (Sony HDR-CX730/HDR-CX200) placed inside or outside of the home enclosures. The cameras ran for about 15–30 minutes to ensure full feeding sessions were captured (as not all animals were feeding at the same speed).

To control for environmental factors that could potentially influence vigilance levels, we implemented a two by two design. We varied 1) the location of the feeding between ‘inside’ (i.e., the inside part of the home enclosure, where animals are normally fed) and ‘outside’ (i.e., the outside part of the home enclosure, accessible via a semi-transparent tube system) and 2) the number of feeding bowls the mash was provided in between one or two bowls while keeping the amount of food constant. The location of the feeding was altered to provide variation in the experienced risk level of the feeding situation. ‘Inside’, where animals were used to being fed, leading to a “lower risk” situation and ‘outside’, where potential predators were visible more frequently, leading to a “higher risk” situation. To account for potential space restrictions, we provided one or two feeding bowls. The animals experienced the four conditions in a semi-randomised order, for a total of 28 sessions collected over 5 weeks in May 2018.

### Data coding

Video coding was done with the software INTERACT (Mangold GmbH, version 18.0.1.10, Arnstorf, Germany). Coding started with the first frame from which the experimenter was no longer visible (location ‘inside’) or the frame when first individual was fully (all four limbs) located outside of the tube that connected the inside to the outside enclosure (location ‘outside’). The following time periods were excluded from coding: 1) when animals were not feeding for more than 4 min. 2) when obvious outside disturbances occurred that would externally induce vigilance, i.e., cats walking past the outside enclosure, other groups of the colony vocalising loudly (as we were interested in the effects of pre-emptive vigilance not reactionary vigilance, see [73]). Sessions were deemed finished when animals stopped eating because all the mash had been eaten, or when they interrupted feeding for more than 4 min. without resuming, or after 10 min. We coded three behaviours namely: vigilance, feeding and out of sight as well as the locations of the animals (inside/outside or on basket) according to the detailed definitions in Table S2. Interobserver reliability was assessed by coding 21 % of all video data by a second rater. We reached interclass correlation coefficients (ICC3) of 0.95 (95% confidence interval, CI [0.86, 0.98]) for vigilance, 1 (95% CI [0.99, 1.00]) for feeding and 0.94 (95% CI [0.85, 0.98]) for the location categories.

### Data preparation for analyses

We used the onsets and offsets of behaviours “vigilance” and “feeding” coded from the videos (according to definitions in Table S2) to determine behavioural bouts of interest. Analysis was restricted to behavioural states when (1) *both* animals were located on the feeding basket (‘together’; to fit the Kuramoto Model and the non-periodic Kuramoto model, see below) or (2) one animal was located *alone* on the feeding basket (‘alone’; for control simulations). Since animals were allowed to move freely in the enclosures, time spent on the basket occurred in bouts. We determined bout length by studying the distribution of breaks in behaviour (i.e., whenever an animal was not displaying vigilance or feeding) and assigned 10s to be a sensible cutoff. These criteria resulted in a total of 53 behavioural bouts from 7 pairs when both individuals were together on the feeding basket (7.6±4.0 mean±s.d bouts per pair). The behavioural bouts were converted to a numerical time series by assigning the following values to the behavioural states: vigilance = 1, feeding = 0. Marmosets also showed behaviours that could not be assigned either to the definitions of vigilance or feeding. These behaviours were considered transition states between vigilance and feeding and were assigned the value of 0.5 (this included behaviours where the line of sight of the individual was not clear, i.e., “out of sight”).

Because marmosets are known to show head movements of angular velocities up to 1000 degrees per second, which equates to 2.78 complete head oscillations per second, we lowpass filtered the time series using a filter of 2.78 Hz (using MATLAB’s ‘lowpass’ function from the ‘signal processing toolbox’). A Hilbert transform was applied to the filtered time series (using MATLAB’s ‘hilbert’ function from the ‘signal processing toolbox’), and a time series of phase angles was obtained.

### Investigating the properties of marmoset vigilance behaviour

We investigated the durations for which an individual showed a particular behaviour before switching to a different behaviour. We modelled these switches in behaviours as Poisson processes and therefore fit exponential functions (using MATLAB’s ‘fitdist’ function from the ‘statistics and machine learning toolbox’) to the distributions of durations. We studied the fit parameter for the two behavioural durations for all individuals, both when they were alone and when they were together. Specifically, we compared the ratio of the fit parameters of the two individuals of a pair for vigilant and feeding durations. We also drew 1000 bootstrap samples from the vigilant and feeding distributions when individuals were ‘alone’ and determined the distribution of the fit parameter ratios (for both vigilance and feeding). We then determined the Z-score and the corresponding p-value of the fit parameter ratio obtained from the ‘together’ condition based on its distribution in the ‘alone’ condition. This represented the probability of obtaining a ratio as extreme as the one from the ‘together’ condition from the ‘alone’ condition by chance. Then, for each individual, we estimated the mean time period of head oscillations to be the sum of the vigilance and feeding fit parameters and compared this time period between the ‘alone’ and ‘together’ conditions. Finally, we checked if the mean time-period of head oscillations is correlated between the individuals of the pair in both the ‘alone’ and ‘together’ conditions.

### Fitting the Kuramoto model

The classic Kuramoto model [64,65] (henceforth ‘Kuramoto model’) for N oscillators is represented by the equation:

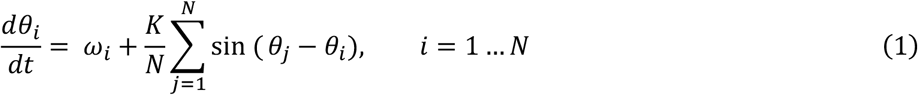

Where *θ*_*i*_ is the phase of the i^th^ oscillator, *ω*_*i*_ is the natural frequency of the i^th^ oscillator, K is the coupling constant and t is time. For the case of two oscillators, it boils down to the following system of equations:

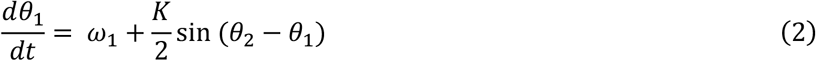

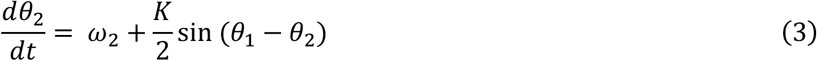

Suppose *ω*_2_ > *ω*_1_. Subtracting equation (2) from (3), we get:

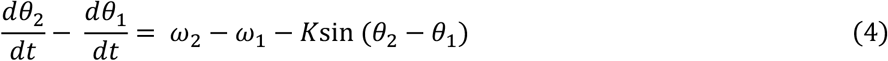

We substitute *θ*_2_ − *θ*_1_ with Φ and *ω*_2_ − *ω*_1_ with Ω to get:

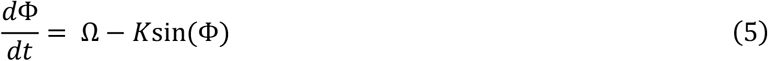

We fit equation (5) to behavioural bouts when both the individuals in a pair were visible by solving it using Euler’s method:

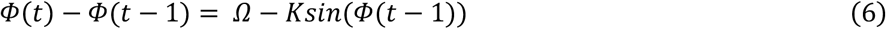

In our case, because the frame rate of the video was 25 fps, the time step size was 0.04s. From the behavioural bouts, we calculated the change in phase difference between the 2 individuals with every frame (Φ(*t*) − Φ(*t* − 1)) and divided it by 0.04 to obtain the value in Hertz. We then regressed Φ(*t*) − Φ(*t* − 1) in Hertz on *sin*(Φ(*t* − 1)) and obtained the slope and the intercept (using MATLAB’s ‘fitlm’ function). *K* was equal to the negative of the slope and Ω to the intercept. A positive *K* indicates the tendency to attain synchrony in-phase and a negative *K* to attain anti-phase synchrony, given the magnitude of *K* is greater than a threshold - the critical coupling constant.

### Determining the critical coupling constant

To determine the critical coupling constant, we visualised the bifurcation diagram (see chapter 3 in [74]) for equation (5) against the control parameter 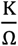 within the limits [-5, 5]. Setting 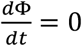 gives:

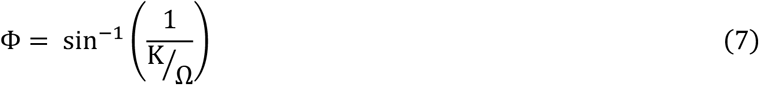

From the bifurcation diagram (figure 1) we see that saddle-node bifurcations occur at 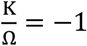 and 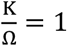 Stable equilibrium states (depicted by solid black curves) exist under conditions 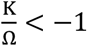 or 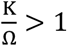. The former condition would cause the system to asymptotically approach anti-phase-like synchrony (as 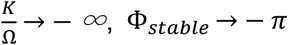) and the latter in-phase-like synchrony (as 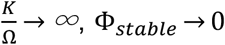). As we were interested in anti-phase synchrony, the critical coupling constant K_c_ would have to follow:

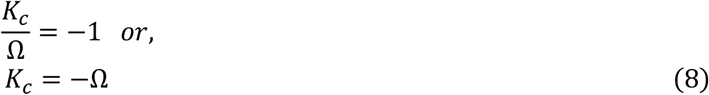

This works, given our initial assumption that *ω*_2_ > *ω*_1_. However, in general,

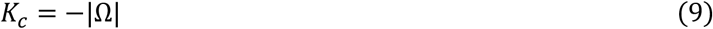

We determined the value of K_c_ for every bout and compared it with the value of K obtained for that bout. Anti-phase synchrony is attained when K<K_c_.

**Figure 1.**
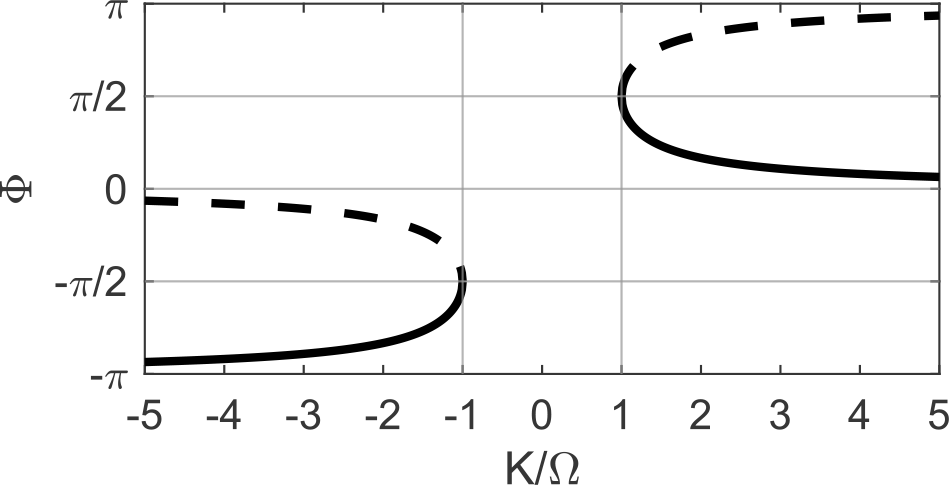
Bifurcation diagram of the Kuramoto model for 2 oscillators. The diagram depicts equilibrium points, i.e., the values of 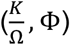 for which 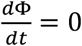. Equilibrium points form continuous curves and are of 2 types: stable, represented by solid curves and unstable, represented by dashed curves. The sign of 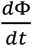 in the close vicinity of the solid curves will cause the system to move towards the curve whereas, in the vicinity of dashed curves, will cause the system to move away from it. Note that the y-axis can be wrapped around a circle as *π* = −*π* on a polar plot, and the same graph can be plotted on the surface of a cylinder.

### Estimating the time to achieve anti-phase synchrony

To estimate the time taken by the marmoset pair to achieve anti-phase synchrony, we first determined the initial phase difference Φ_*i*_ between the two individuals from the time series of phase angles (see paragraph on data preparation for analyses). We assumed that at time t=0, the phase difference between the individuals Φ = Φ and at t=T, they reach anti-phase synchrony, i.e., 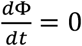 or, from equation 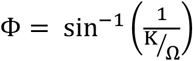. To calculate T, we integrate equation (5) as follows:

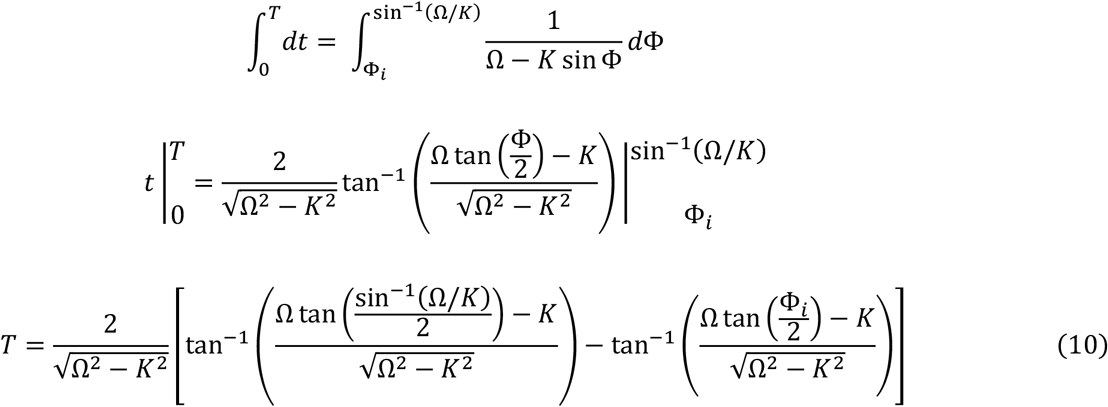

From the values of Φ_*i*_, Ω and *K* obtained for every bout, we calculated the time taken to reach anti-phase synchrony using equation (10).

### Control simulations

We simulated 100 behavioural bouts for every marmoset pair based on the behaviour of each of the individuals of the pair when they were situated alone on the feeding basket. The duration of every simulated bout was set to a random number obtained from an exponential distribution fit to the histogram of bout durations of the pair of interest. Then, for each individual in the pair, we determined the distribution of durations between a feeding-vigilant transition and the subsequent vigilant-feeding transition (‘vigilant durations’) and, similarly, obtained the ‘feeding durations’. Next, we simulated behavioural bouts for each individual of the pair separately, considering only its vigilant and feeding durations, and no coupling between them. For each pair, we initiated 25 simulations from the vigilant-vigilant (= state of individual 1 - state of individual 2) state, 25 from vigilant-feeding, 25 from feeding-vigilant and 25 from feeding-feeding. Once initiated, a random number from the exponential distribution fit to that individual’s vigilant and feeding durations determined at what point the individual switched its state. For example, let us say we want to simulate data for individuals A and B starting with the state feeding-vigilance. So, the value of behaviour of A at time 0 is 0 and for B at time 0 is 1. We pick a random number from the exponential distribution fit to A’s feeding durations; let us say it is 6.5. Then, the value for the behaviour of A remains 0 until time=6.5s and then switches to 1. We now pick a random number from the exponential distribution fit to A’s vigilant durations; let us say it is 3.3. Then A retains the value of 1 from time=6.5s until time=(6.5+3.3)=9.9s and then switches to 0. This continues until the bout duration (determined previously; see above) is reached. The same is done for B.

All behavioural bouts obtained were lowpass filtered with a filter of 2.78 Hz (using MATLAB’s ‘lowpass’ function from the ‘signal processing toolbox’), and Hilbert transform applied (using MATLAB’s ‘hilbert’ function from the ‘signal processing toolbox’) to obtain time series of phase angles. The Kuramoto model was fit to the simulated bouts, and coupling constant and critical coupling constant values were obtained using the same procedure described previously.

### The non-periodic Kuramoto model

The Kuramoto model assumes that all oscillators are inherently periodic. However, Sarfati et al. [67] recently showed that fireflies that do not inherently periodically flash can synchronise their flashing. Inspired by this phenomenon and the fact that we do not expect individual marmosets to demonstrate perfect periodicity in head oscillations, we came up with a modified form of the Kuramoto model, which does not assume that the oscillators are inherently periodic. We propose the equations of this non-periodic Kuramoto model for the marmoset pairs in our data by modifying equations (2) and (3) as follows:

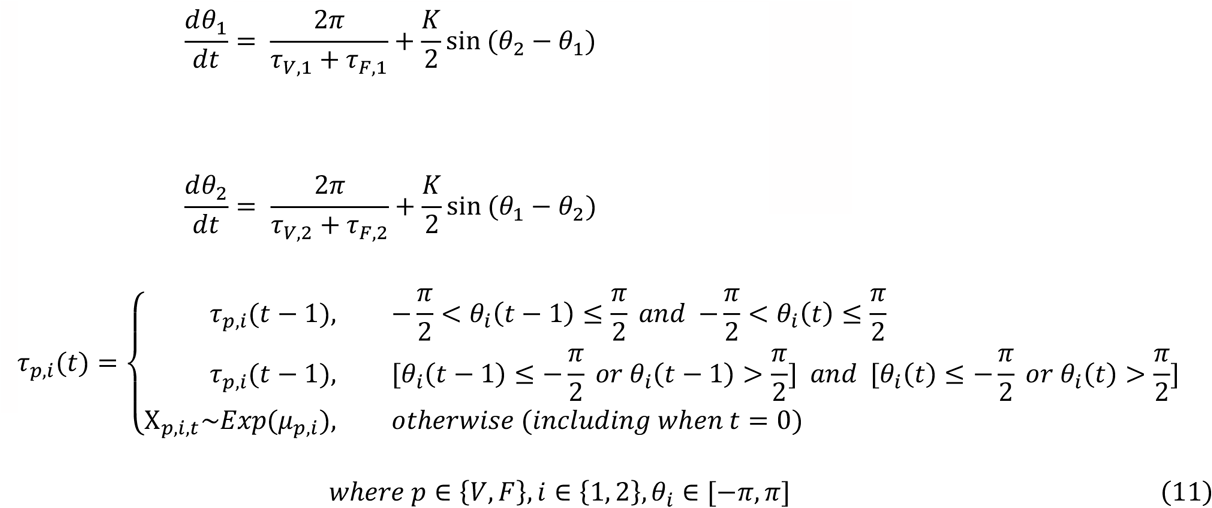

Here, subscript V stands for the vigilant state and F for the feeding state. *μ*_*p,i*_ is the mean of the exponential distribution fit to durations of state p for individual i, and X_*p,i,t*_ is a random number drawn from that exponential distribution at time t. The random variable τ_*p,i*_, which represents the time period for which individual i would remain in state p, makes the non-periodic Kuramoto model different from the classic model. In the non-periodic model, the natural frequency of an individual resets to a random number obtained from the empirical data whenever the individual undergoes a transition in the state. Transitions occur at 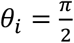 and 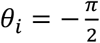.

### Fitting the non-periodic Kuramoto model to the data

For all individuals in the data, we determined the mean of the exponential distribution fit to vigilant durations (*μ*_*V,i*_) and feeding durations (*μ*_*F,i*_). Then, for every bout in the experimental data, we determined the duration and the initial phases of the individuals (*θ*_1_(0) and *θ*_2_(0)). Using these initial phases as the starting point, we performed numerical simulations of equation (11) employing Euler’s method and performing a parameter sweep of K between values -5 Hz and 0 Hz and step size of 0.1 Hz (equating to 50 values of K). For each value of K, we performed 10 numerical simulations with different random number seeds, which slightly varied the output due to the behaviour of the random variable τ_*p,i*_. Therefore, for every behavioural bout in experimental data, we performed 500 non-periodic Kuramoto model simulations. For every simulation, we obtained the time series of phase differences (*θ*_2_ − *θ*_1_). Then, for each value of K we determined the mean of the angular distance between each of the 10 simulated time series of phase differences and the time series of phase differences from the actual behavioural bout. By performing the parameter sweep of K, we determined the value of K for which the angular distance was the minimum. This K-value provided the best fit of the non-periodic Kuramoto model to the data and was, therefore the estimated coupling constant for the behavioural bout.

### Statistical analysis

We used non-parametric tests to test our hypotheses for two reasons: (1) our sample consisted of n=14 individuals, but most tests compared pairs which were 7 and (2) in many cases, we were dealing with ratios and exponential distributions where we couldn’t assume that the distribution underlying the outcome variable was normal. For comparing 2 groups, we applied Wilcoxon signed-rank tests (using MATLAB’s ‘signrank’ function from the ‘statistics and machine learning toolbox’) and for more than 2 groups, we used Friedman’s test (on RStudio using ‘FriedmanTest’ from the package ‘PMCMRPlus’) as the global statistical test followed by post-hoc Nemenyi test (on RStudio using ‘frdAllPairsNemenyiTest’ from ‘PMCMRPlus’) to find which groups differed significantly if the Friedman test detected a significant difference. To compare linear regression lines, we performed non-parametric analysis of covariance (non-parametric ANCOVA on RStudio using ‘sm.ancova’ from the ‘sm’ package). Further, to check for effects of the nature of the session (inside with 1 bowl/inside with 2 bowls/outside with 1 bowl/outside with 2 bowls), we compared linear regression lines of the 4 types of sessions for the actual bouts again using non-parametric ANCOVA.

To investigate the effect of the location of the marmosets (inside/outside) and the number of bowls (1/2) on the coupling constant, we first checked if the coupling constant values obtained from bouts followed a normal distribution using a qq-plot and Kolmogorov-Smirnov test. Because the values followed a normal distribution (p<0.001, Kolmogorov-Smirnov test for normality), we fit a Gaussian linear mixed-effect model with location, number of bowls and the interaction between them as fixed effects and the group as the random intercept. Further, using ANOVA, we compared our model to the null model.

### Overview of analyses

The analysis pipeline is summarised in figure 2.

**Figure 2.**
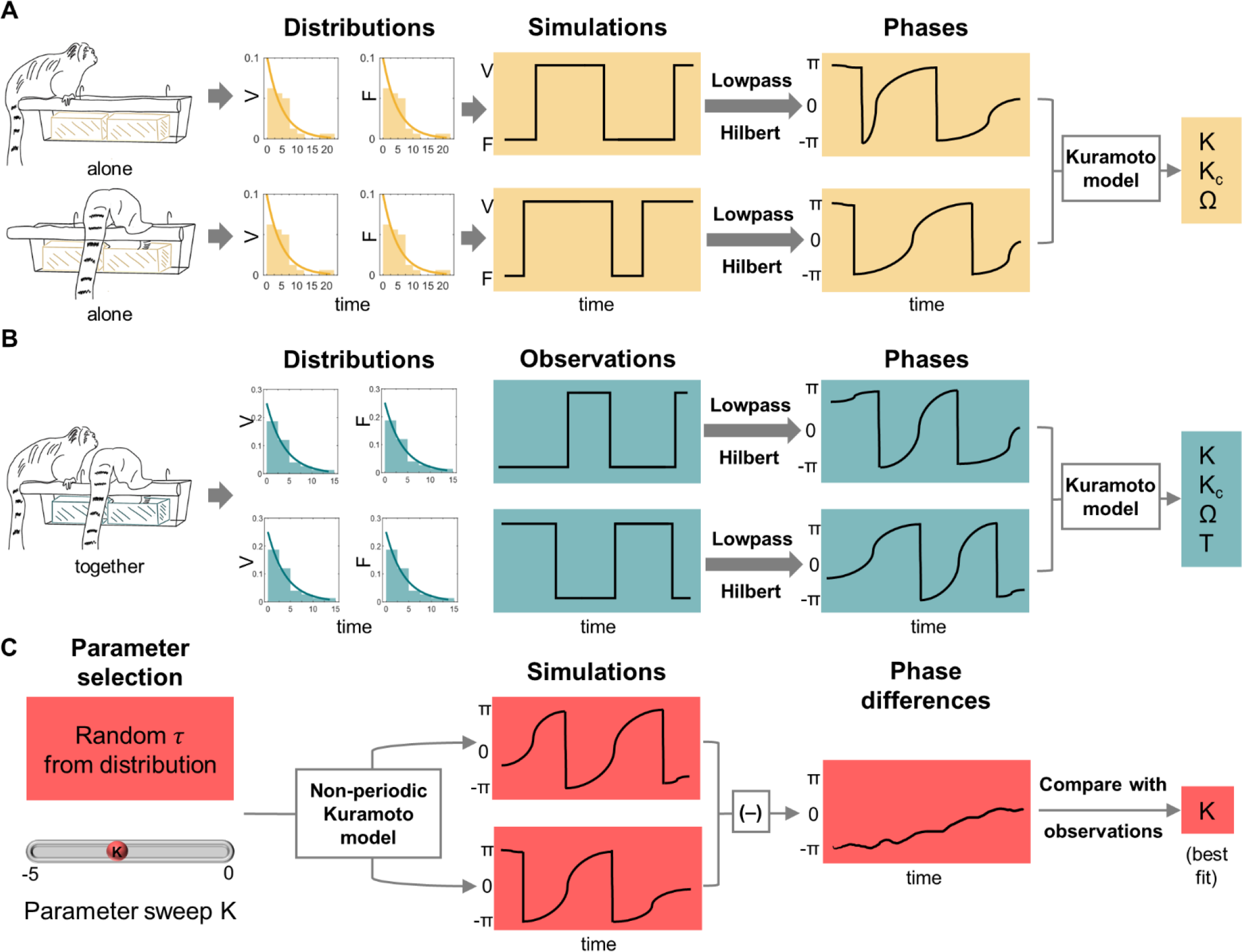
Overview of analyses. **(A) The pipeline for analysing control bouts**. Individuals were observed when they were feeding alone, and their vigilant and feeding distributions were obtained (an individual sat on the basket and ate from the yellow bowl). The distributions were used to simulate 100 behavioural bouts per pair. These bouts underwent lowpass filtering and Hilbert transform, and the Kuromoto model was fit to the phases to obtain synchronisation parameters. **(B) The pipeline for analysing actual bouts using the classic Kuramoto model**. Individuals were observed when they were feeding together, and their vigilant and feeding distributions were obtained. Additionally, the time-series of when each individual was vigilant or feeding was obtained. These time-series (bouts) underwent lowpass filtering and Hilbert transform and Kuromoto model was applied on the phases to obtain synchronisation parameters as in the control condition. **(C) Non-periodic Kuramoto model**. The distributions of vigilant and feeding durations were provided as inputs to the non-periodic Kuramoto model and a parameter sweep of coupling constants done to simulate 2 time-series of phases. From this, the time-series of phase difference was calculated and its angular distance from the time-series obtained from a behavioural bout of the ‘together’ condition was determined. The coupling constant value, which gave the smallest angular distance (provided the ‘best fit’) was the estimated coupling constant for that behavioural bout. Note that even though monkeys are depicted to be provided with one food bowl when alone and two food bowls when together, both one bowl and two bowl conditions were experienced by all animals, Abbreviations: V=vigilance, F=feeding, K=coupling constant, K_c_=critical coupling constant, Ω=difference in natural frequencies, T=time to attain anti-phase synchrony, and τ=time-period for which an individual would remain in a particular behavioural state.

## Results

### Properties of marmoset vigilance behaviour

The distributions of durations for which an individual displayed a behaviour before switching to a different behaviour were right-skewed (see figure 3A for distributions of an example pair). We modelled these switches in behaviours as Poisson processes and therefore fit exponential functions to the distributions of durations. We found the fit parameter *μ* for vigilance distributions to be more similar between individuals of a pair (ratio between the individuals significantly higher and closer to 1) when they were together compared to when they were alone (figure 3B, n=7 pairs, signed-rank=1, p=0.03, two-sided Wilcoxon signed-rank test). Values closer to 1 suggest synchrony. Moreover, for four pairs, the probability of obtaining a ratio as extreme as in the ‘together’ condition from the ‘alone’ distributions by chance was <0.05. Significant differences were not found for feeding distributions (figure 3C, n=7 pairs, signed-rank=4, p=0.11, two-sided Wilcoxon signed-rank test). However, even in this case, the probability of obtaining a ratio as extreme as in the ‘together’ condition from the ‘alone’ distributions by chance was <0.05 for three pairs.

**Figure 3.**
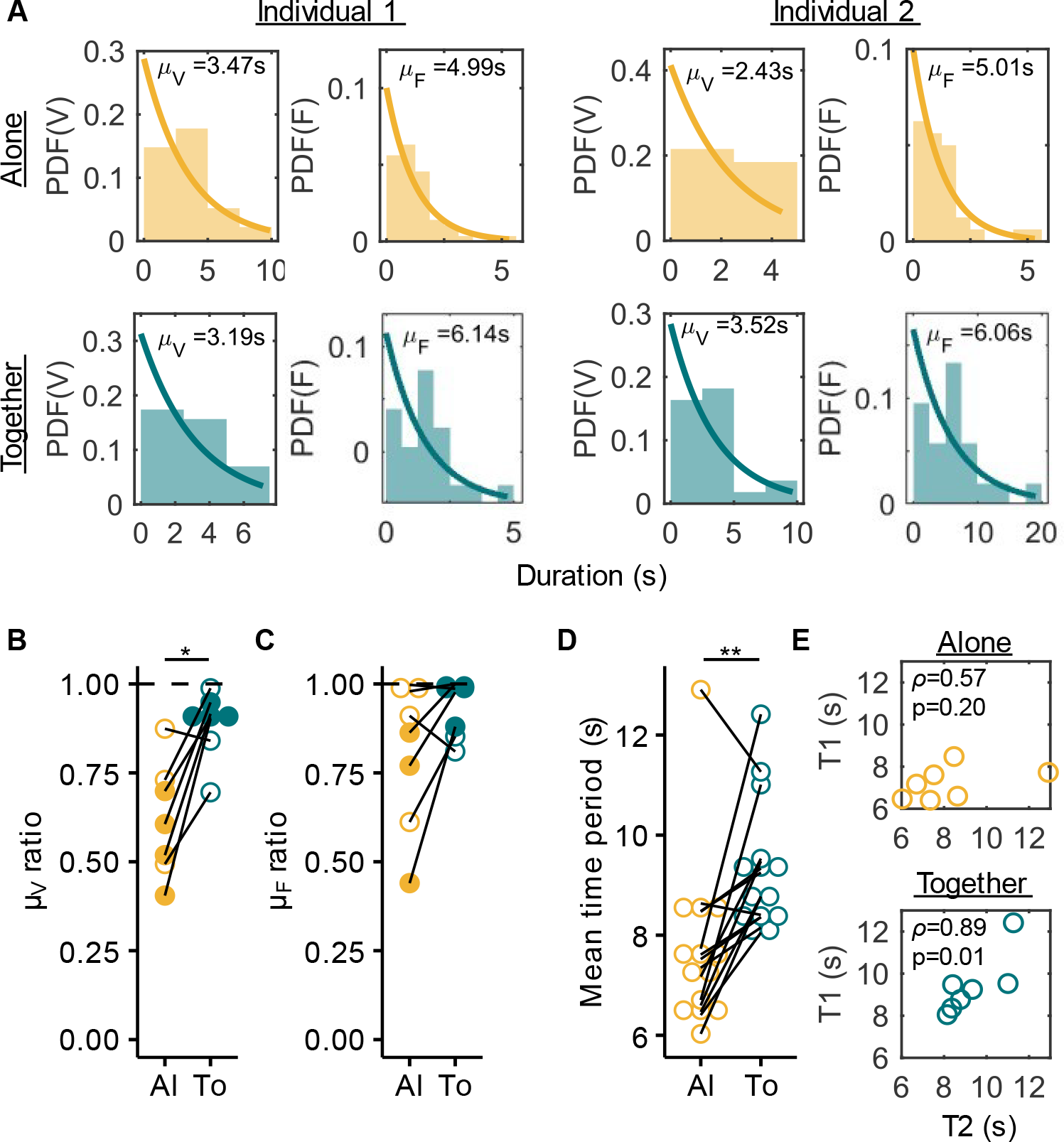
Properties of marmoset vigilance behavior. **(A)** Distribution of behavioural durations for vigilance (V) and feeding (F) behaviours for an example pair in the data. The two left columns correspond to one individual and the two right columns to the other individual of the pair. The top row depicts the distributions when the individuals were alone and the bottom when they were together. Each bin of the histogram is 2.5s. Exponential fits to the probability density functions (PDFs) are shown with their corresponding parameter values (mean *μ*). **(B, C)** Plots comparing the ratio of mean vigilance durations (*μ*_*V*_ ratio, **B**) and mean feeding durations (*μ*_*F*_ ratio, **C**) of the two individuals of the pairs (n=7 pairs) between the alone (‘Al’) and together (‘To’) conditions. Each point is a pair and the same pairs are connected by lines. Values closer to 1 mean that the distributions of durations for the two individuals are similar, a sign of synchrony. Furthermore, if the probability of obtaining the together value by chance from the alone distributions was <0.05 (see methods), filled circles are shown instead of open circles. *p<0.05, two-sided Wilcoxon signed-rank test. **(D)** Plot compares the mean time period (*μ*_*F*_ + *μ*_*V*_) of n=14 individuals between the alone and together conditions. Each point is an individual, and the same individuals are connected with lines. **p<0.01, two-sided Wilcoxon signed-rank test. **(E)** Correlation between the mean time periods (*μ*_*F*_ + *μ*_*V*_) of the individuals of the pair in alone (top) and together (bottom) conditions. Spearman’s *ρ* and the corresponding p-values are shown.

While ratios of fit parameters alluded towards synchrony in the ‘together’ condition, they cannot indicate the nature of synchrony, i.e., whether the marmosets showed in-phase or anti-phase synchrony. For this, we compared the estimated mean time-period of head oscillations (*μ*_*F*_ + *μ*_*V*_) between the two conditions and found it to be significantly higher when the individuals were together (figure 3D, n=14 individuals, signed-rank=8, p=0.0031, two-sided Wilcoxon signed-rank test). This effect was mostly driven by a significant increase in feeding durations of the individuals (figure S1). It indicated that the individuals were most likely in anti-phase synchrony when together. More evidence for synchrony came from the fact that the mean time period of head oscillations between the individuals of a pair was positively correlated when they were together (figure 3E bottom panel, n=7 pairs, Spearman’s coefficient=0.89, p=0.01) and not when they were alone (figure 3E top panel, n=7 pairs, Spearman’s coefficient=0.57, p=0.2).

### Formulating the models

The Kuramoto model assumes that the individual oscillators are periodic. The Kuramoto model for N oscillators is given by:

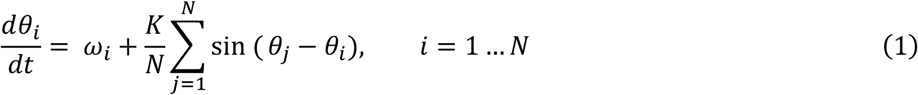

Where *θ*_*i*_ is the phase of the i^th^ oscillator, *ω*_*i*_ is the natural frequency of the i^th^ oscillator, K is the coupling constant and t is time.

From equation (1), we propose an extension to the Kuramoto model for non-periodic oscillators. This extension is similar to the extension to the integrate-and-fire model for non-periodic fireflies proposed by Sarfati et al. [67] Let there be N non-periodic oscillators, each going through a sequence of M states to complete one oscillation and the state of the i^th^ oscillator at time t be represented by Ψ_*i*_. Suppose the probability of transitioning from one state to all other states except the next one in the sequence is 0. Let the duration of the i^th^ oscillator to remain in the p^th^ state before transitioning to the next one be denoted by T_*p,i*_, be independent of its duration to remain in any other state, and follow the distribution *g*(T_*p,i*_). Then equation (1) can be modified to:

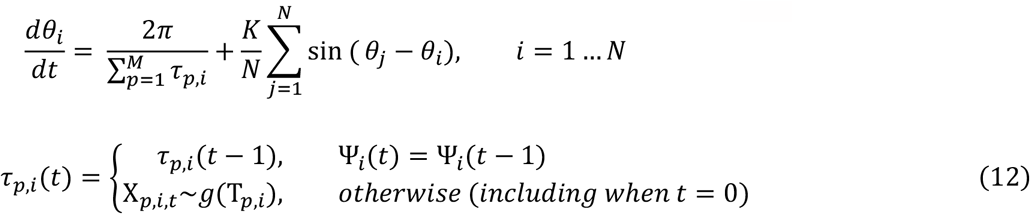

where X_*p,i,t*_ is a random number drawn from the distribution *g*(T_*p,i*_) at time t and τ_*p,i*_ is a random variable who’s value changes whenever the state of the oscillator changes.

Simulations of equation (12) for the case of marmosets where two non-periodic oscillators switch between two states (see methods) show that non-periodic oscillators can display both in-phase and anti-phase synchrony depending on the value of the coupling constant (figure 4).

**Figure 4.**
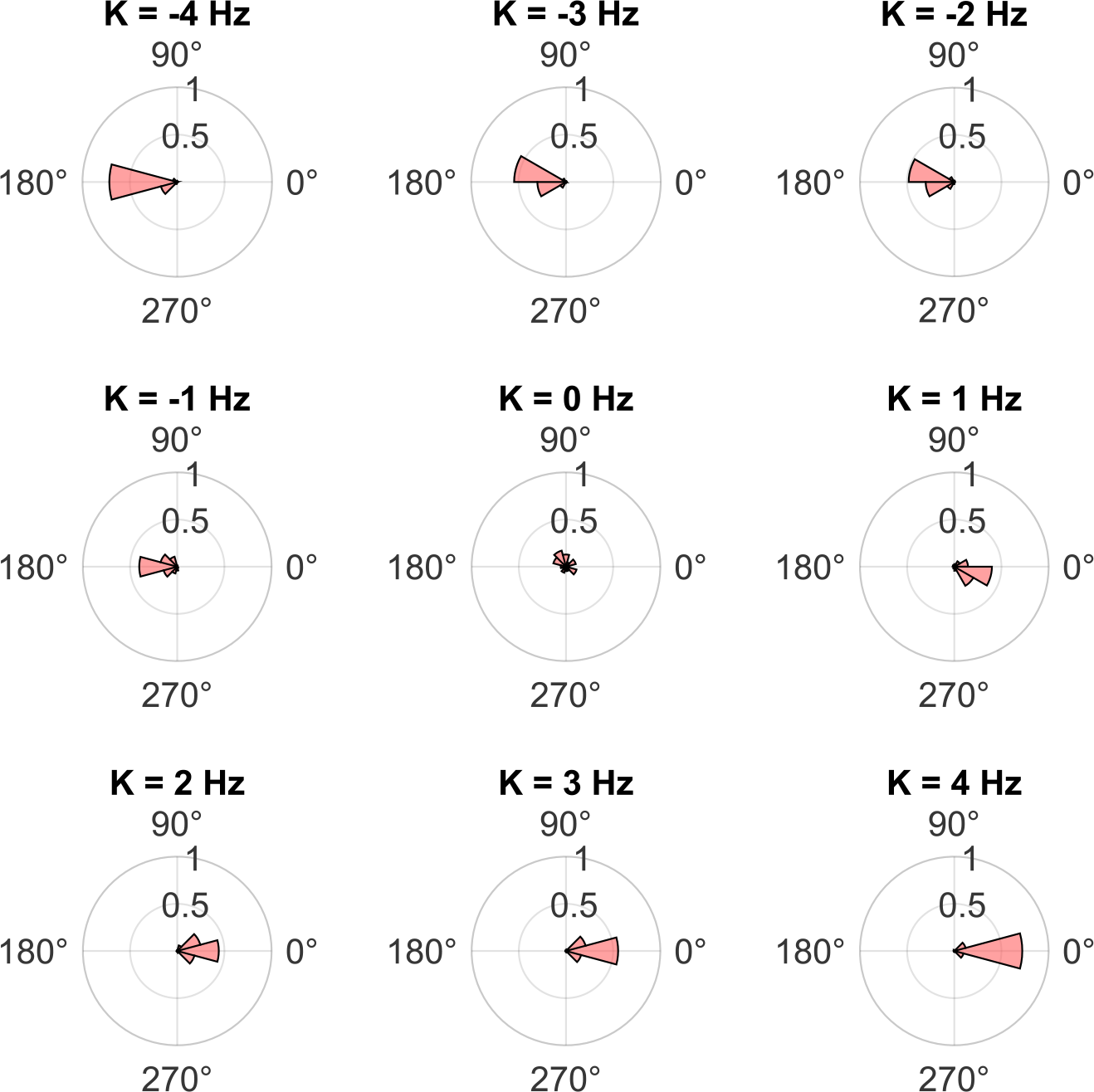
Synchrony in the non-periodic Kuramoto model. Example polar probability histograms of the phase differences between 2 non-periodic Kuramoto oscillators (see equation 11) for different values of the coupling constant (K). The parameter values for the model simulations were the average of values in the data (*μ*_*F*,1_=5.06s, *μ*_*V*,1_=3.17s, *μ*_*F*,2_=4.04s, *μ*_*V*,2_=3.10s, duration of simulation=53.32s=1333 time steps), and the initial phase difference was 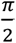.

### Results from model fits

Out of the 53 behavioural bouts from 7 marmoset pairs in the data, the Kuramoto model could only be fit to 47 bouts, the remaining 7 bouts being either too short in duration (<10s) or consisting of too few head oscillations (<2 for both individuals) (see figure S2 for the histogram of bout durations before and after discarding the 7 bouts). Along with those seven bouts, we also discarded all bouts that were less than 10s long or in which both individuals underwent less than 2 complete head oscillations. This provided us with 38 bouts to analyse.

The coupling constant (K) from the Kuramoto model fit was more negative than the critical coupling constant (K_c_) for 35 out of the 38 bouts (i.e., 92.1% of all bouts, figure 5A, B). Based on this model, the average time required for each pair to reach anti-phase synchrony was 6.84±2.27s (mean±sd). Overall, the fraction of bouts for which K<Kc was significantly higher for actual pairs than control pairs (figure 5C, signed-rank=28, p=0.016, two-sided Wilcoxon signed-rank test). The K-values of the control pairs, actual pairs with the classic Kuramoto model fit, and actual pairs with the non-periodic Kuramoto model fit were significantly different (figure 5D, *X*^*2*^=14, df=2, p<0.001, Friedman test). Post-hoc analysis showed that control pairs and actual pairs with the non-periodic Kuramoto model fit differed significantly (control pairs – non-periodic Kuramoto: p<0.001, control pairs – actual pairs (Kuramoto): p=0.147, actual pairs (Kuramoto) – non-periodic Kuramoto: p=0.147, post-hoc Nemenyi test). Further, there was no effect of the location or the number of bowls in themselves, but we did find a significant effect of their interaction on the K-value (table S3). This was driven by a more negative K-value when the marmosets were outside and provided with 2 bowls (figure S3). The Gaussian linear mixed-effect model was significantly different from the null model (log-ratio=11.18, p=0.01).

**Figure 5.**
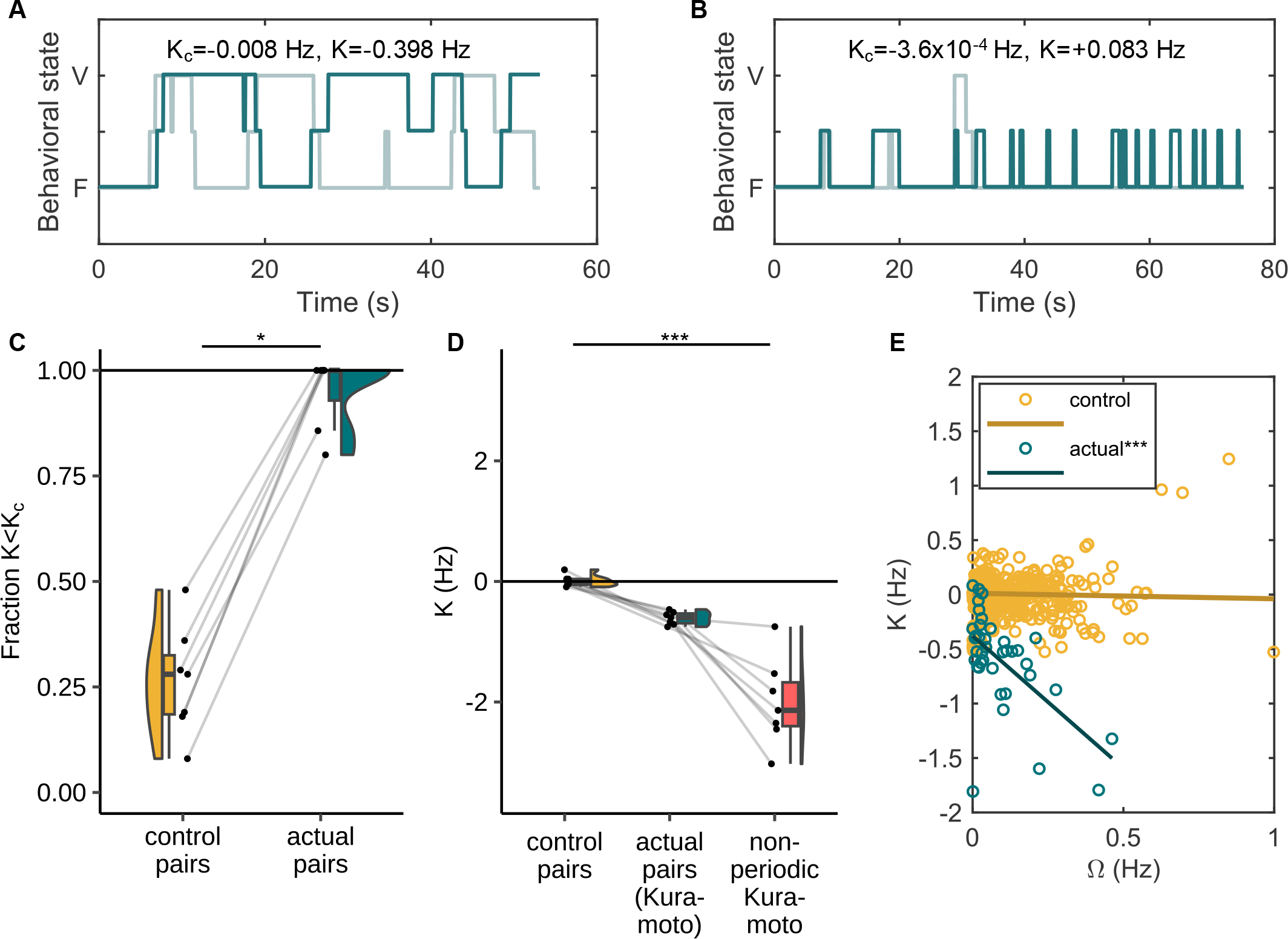
Synchrony in control and actual pairs. **(A, B)** Example behavioural bouts from actual pairs. Plots show time-series of behaviours of the two individuals of a pair when they were together. Each curve (dark or light teal) corresponds to one individual. ‘V’ stands for vigilant state and ‘F’ for feeding. Here, K_c_ and K are obtained from the classic Kuramoto model fit. 92.1% of the analysed data (from actual pairs) consisted of bouts such as in (A) where K<K_c_ and only 7.9% consisted of bouts such as in (B) where K>K_c_. **(C)** Fraction of behavioural bouts out of total for which the coupling constant was less (more negative) than the critical coupling constant for control and actual pairs (n=7). *p<0.05, two-sided Wilcoxon signed-rank test. **(D)** Coupling constant (K) in control pairs, actual pairs with the classic Kuramoto model fit and actual pairs with non-periodic Kuramoto model fit. Each point is a mean of K-values of all bouts of that pair (n=7 pairs). ***p<0.001, post-hoc Nemenyi test. **(E)** Relationship between the critical coupling constant and the difference in natural frequencies of the two individuals for control bouts (simulated, n=700) and actual bouts (n=38). Lines show linear regression fits to control and actual bouts. ***p<0.001, non-parametric ANCOVA.

Across the 38 bouts, the K-value from the classic Kuramoto model negatively covaried with the difference in natural frequencies (Ω) between the individuals of the pair (figure 5E, slope of linear regression=-2.43, y-intercept=-0.38 Hz, RMSE=0.36 Hz). Linear regression analysis of K-values of control bouts (n=700) and how they varied with Ω gave negligible slope and intercept (figure 5E, slope=-0.05, y-intercept=0.01 Hz, RMSE=0.49 Hz). Non-parametric analysis of covariance (non-parametric ANCOVA) showed that the two regression lines were unequal (error standard deviation=0.22, p<0.001). Further, there was no effect of the location (inside/outside) or the number of bowls (one/two) on the regression lines (non-parametric ANCOVA, error standard deviation=0.32, p=0.233).

## Discussion

In this study, we show that pairs of marmosets coordinate vigilance by displaying anti-phase synchrony in their behaviours despite each individual being non-periodic. More interestingly, we find that coupling between individuals is flexible and there is increased coupling when the individuals are inherently more asynchronous. Below we discuss a plausible mechanism for the above phenomena, its implications for marmoset behavioural ecology, and the wider applicability of our mathematical framework for the study of synchronisation in nature.

### Implications for marmoset behavioural ecology

Marmosets are not perfectly periodic oscillators when it comes to displaying vigilance and feeding behaviours. When alone, the distributions of durations for which each individual marmoset was vigilant or feeding before changing its behaviour followed an exponential distribution (see figure 3A for an example). This property made it ideal to model the behaviour as a Poisson process. It suggests that marmosets have an underlying rate of change of behaviour (vigilance/feeding); however, each transition in behaviour is more or less independent of the other.

Marmosets displayed changes in behaviours when they were together. The distributions of durations of being vigilant between the two individuals of a pair became more similar when together compared to when alone (figure 3B), a sign of synchrony. The feeding distributions also became more similar in the ‘together’ condition, even though this effect was not significant (figure 3C). Overall, the mean time period of oscillations (vigilance and feeding) of the two individuals of a pair were positively correlated when together, which was not the case when alone (figure 3E). This suggests that becoming similar to each other in vigilance durations along with a moderate level of similarity in feeding durations was sufficient to display overall synchrony. Moreover, marmosets spent longer in a behavioural state before changing their behaviour when together, effectively slowing down head oscillations compared to when alone (figure 3D). Such slowing down of oscillations has been proposed to be a phase-delay mechanism of attaining anti-phase synchrony and has been described in case of alternating singing in several insect and frog species [1,19]. In katydid pairs for example, it has been proposed that a chirp from one individual delays the chirping of the other individual by a tiny bit (less than one cycle), thus inducing a phase-delay [75]. This applies symmetrically to both individuals and over time the individuals display anti-phase synchrony, with the frequency of chirping of each individual reduced.

Based on the slowing down of head oscillations, followed by the confirmation of negative coupling from the Kuramoto model fit to the data (figure 5D), a mechanistic explanation could be that when a marmoset sees another individual change its behavioural state (feeding/vigilant), it remains in its current state for slightly longer than usual. This applies symmetrically to both individuals and the accumulation of phase-delays ultimately leads to anti-phase synchrony. As to why marmosets may remain in a state for prolonged duration when together, an ethological explanation could be that a marmoset may feel ‘safer’ in presence of the other individual, hence prolonging the time it would feed. The other individual, if vigilant, would now have to accommodate to this by prolonging its vigilance state, altogether triggering a chain of phase-delays that would ultimately lead to anti-phase synchrony. This ethological hypothesis is supported by the fact that the increase in feeding durations was the primary contributor to the slowing down of head oscillations (figure S1), whereas accommodating to each other’s vigilance durations was the primary contributor to developing and maintaining synchrony (figure 3B, C).

A negative coupling alone will not lead to anti-phase synchrony. The coupling must be strong enough to overcome the mutual differences in natural frequencies of the two individuals that prevent them from synchronising. By fitting the Kuramoto model, we found that in 92.1% of the analysed bouts, the coupling between individuals was strong enough to overcome the critical threshold to attain anti-phase synchrony, almost four-fold higher than what was expected by chance (figure 5C), hence confirming prediction 1 (see introduction). Moreover, the coupling constant estimates from the non-periodic Kuramoto model fit were more negative than the Kuramoto model fit. This is intuitive, as the marmosets would have to couple more strongly to display the same temporal progression of synchrony if they were not inherently periodic. Furthermore, the coupling was stronger when the marmosets were outside and had access to two bowls (figure S3). The stronger coupling when outside can be attributed to the increase in perceived risk that could have motivated them to attain anti-phase synchrony sooner. The coupling being stronger in the presence of two bowls tells us that the marmosets were not merely forced to show anti-phase synchrony by the provision of one bowl only. If it was the case that the lack of feeding space (in the case of 1 bowl) was the motivation behind coordinating vigilance, then we would have expected stronger coupling in the presence of 1 bowl, which was not the case.

In line with prediction 2 (see introduction), marmosets displayed a high degree of flexibility in coupling. The coupling strength ranged from -1.81 Hz to +0.08 Hz. This variability was not random; the more the difference was between two pair mates in natural frequencies as estimated by the Kuramoto model, the stronger and more negative was the coupling strength (figure 5E). Needless to say, this is not expected to happen by chance (see control bouts in figure 5E) as differences in natural frequencies and coupling strengths are two independent parameters of coupled oscillator systems. Nor does the simple mechanistic description (see above) explain such a trend. Marmosets seem to put in more effort to synchronise with each other (by coupling more strongly) when they begin in a more desynchronised state, confirming prediction 3 (see introduction). This makes the process cognitively more demanding as it would require the marmosets to estimate the amount of asynchrony i.e., the differences in rhythms, and mutually compensate for larger differences in rhythms with a stronger coupling. However, currently, we do not have enough understanding of the cognitive aspect of synchronisation in marmosets to claim if this a conscious process or otherwise. What we do know is that similar patterns are found when marmosets accommodate to their partners in other modalities as well. For example, marmosets have been shown to converge to their partner in vocal space over time [70,72] (called “vocal accommodation” [69]) and it has been observed that the more the individuals differ in vocal properties initially, the more they converge. The simplest mechanistic explanation obtained from mathematical modelling of the phenomenon suggests the presence of a dynamic vocal learning mechanism [71], which is again cognitively more demanding than simple forms of learning. Overall, it is clear that mathematical modelling is a powerful tool that could help us tap into the flexibility and control that animals might have over synchrony parameters and, ultimately, the cognitive aspects of synchrony. Marmosets with their ability to flexibly synchronise with other individuals in various modalities such as in motor coordination [76] (for evidence of motor coordination in another other callitrichid species see [77]), vocal turn taking [12,13,78,79] and peripheral oxytocin levels [80] make for an ideal animal model to study this phenomenon. [76,81].

### Implications for studying synchrony in nature

Weakly coupled oscillators are prominent motifs in the animal world, exemplified by gatherings of fireflies flashing together [4], crickets producing synchronised chirps [19], and animals calling antiphonally [12,14,15]. These systems take a finite time to attain synchrony if they begin in an unsynchronised state. Therefore, if an observer observes the system before it attains synchrony, they will not see the system to be synchronized but can only note that the system is tending towards synchrony. In such cases, looking at mean phase coherence, distribution of phase differences, or overall correlation may not necessarily show this trend. By fitting the Kuramoto model to the system, one can detect such a trend towards synchrony. Moreover, for N=2 oscillators case, we show that it is possible to estimate the time required for the system to attain synchrony (equation 10), which allows for assessing the biological relevance for the system under study. For example, if it would take 30 minutes instead of 6 seconds for the marmosets to attain anti-phase synchrony, this could hardly be a good predator avoidance strategy during feeding bouts, even though the coupling constant from the Kuramoto model fit in such a case would be more negative than the critical coupling constant.

A drawback of the Kuramoto model is that it assumes that each oscillator it is modelling is inherently periodic. However, in biology, we seldom come across perfectly periodic oscillators. The Kuramoto model can indeed tolerate some level of variability in natural frequencies, but when the distributions are not narrow, as in the case of *Photinus carolinus* fireflies’ inter-flash intervals [82] or vigilance and feeding durations of marmosets, modifications to previous models are required. Akin to Sarfati et al.’s extension to the integrate-and-fire model [67], our extension to the Kuramoto model allows users to implement it even when individual oscillators are not perfectly periodic and makes the model applicable to a much wider spectrum of biological phenomena. While the non-periodic model fit provides a more realistic estimate of the coupling strength, estimating other parameters, such as the critical coupling constant or time required to attain synchrony, is not straightforward. Additionally, unlike the classic Kuramoto model, one cannot ‘fit’ the non-periodic Kuramoto to the data in the traditional sense. Estimating the coupling strength requires simulating the model several times by performing a parameter sweep. We, therefore, suggest using the classic and non-periodic models as complementary approaches, as, along with providing critical coupling constant and time to attain synchrony estimates, the classic Kuramoto model also helps narrow down the range of coupling constant values to perform the parameter sweep. In closing, we emphasise the utility of mathematical modelling and its power of abstraction to better understand complex biological processes. We hope that by adding to the long list of demonstrations of the capabilities of mathematical abstraction, we inspire more and more biologists to harness this tool to advance and enrich their field.

## Ethics

All experiments were carried out in accordance with the Swiss legislation and licensed by the Kantonales Veterinäramt Zürich (licence number: ZH223/16; degree of severity: 0).

## Authors’ contributions

N.P., R.K.B. and J.M.B. conceptualised the study. R.K.B. collected the behavioural data. N.P. developed the models and fit them to the data. N.P. analysed the data with inputs from R.K.B.. N.P. and R.K.B wrote the original draft. J.M.B. edited the draft, acquired funding and provided supervision throughout the project.

## Acknowledgements

We thank A. Götschi for data coding, A. Godard for help with interobserver reliabilities and G. Bazzell for animal caretaking as well as support during data collection.

## SUPPLEMENTARY MATERIALS

**Table S1.**
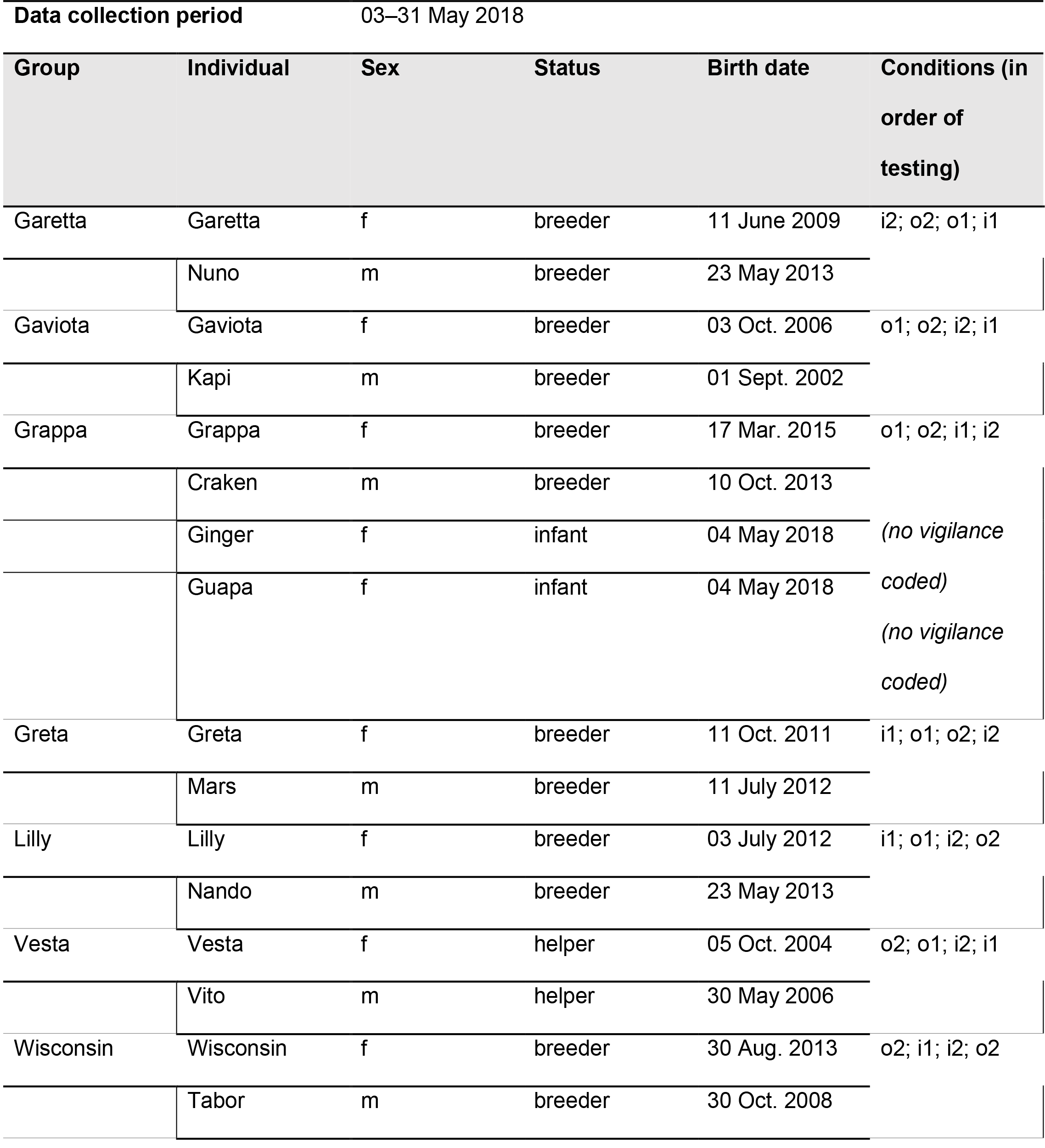
Individuals included in study. List of all individuals involved in the study detailing their age, status and time as well as order of testing. i = inside; o = outside; 1 = one food bowl; 2 = two food bowls

**Table S2.**
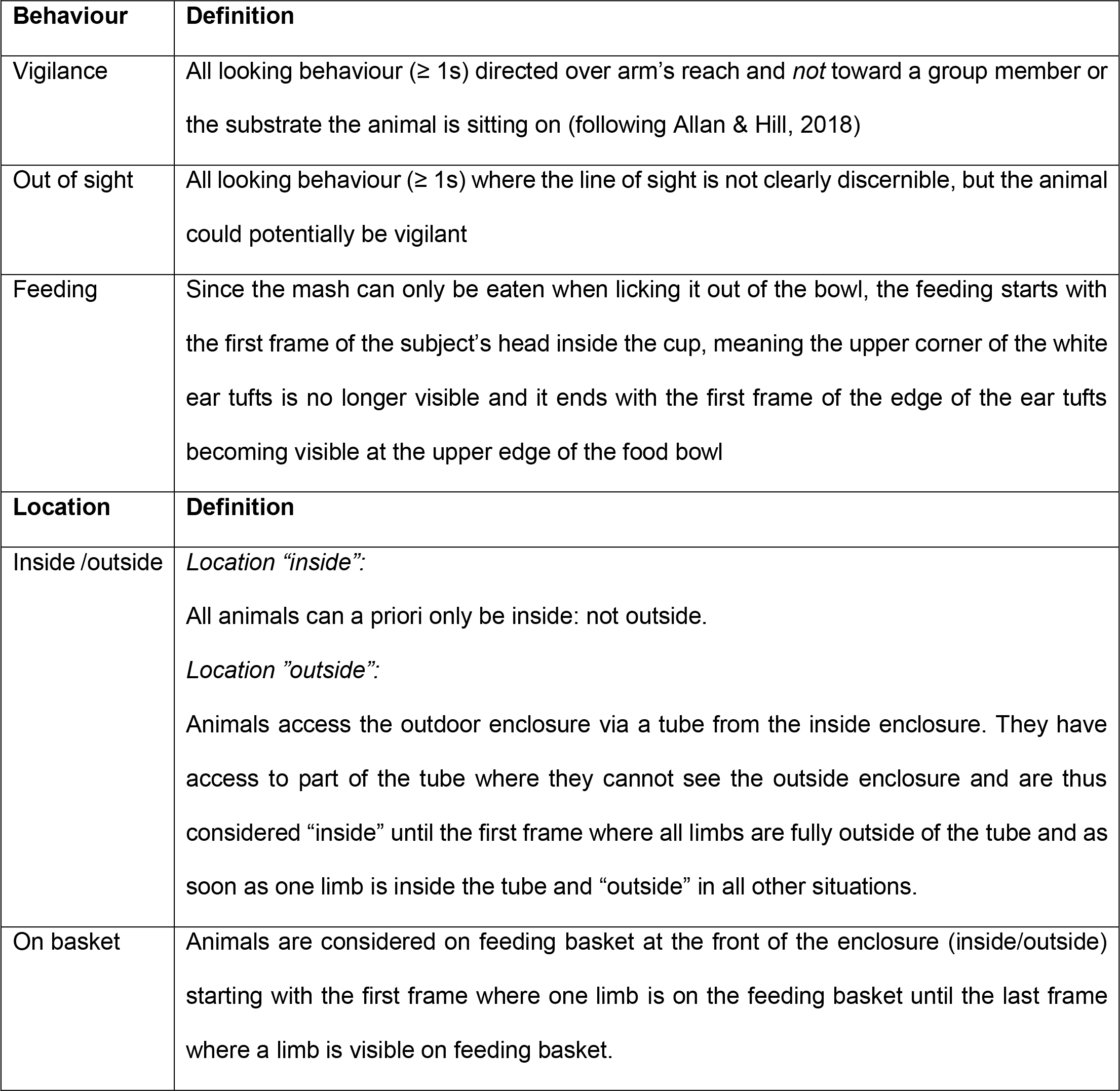
Ethogram with definitions for all behaviours/categories coded from video material. Table taken from Brügger et al. 2023.

**Table S3.**
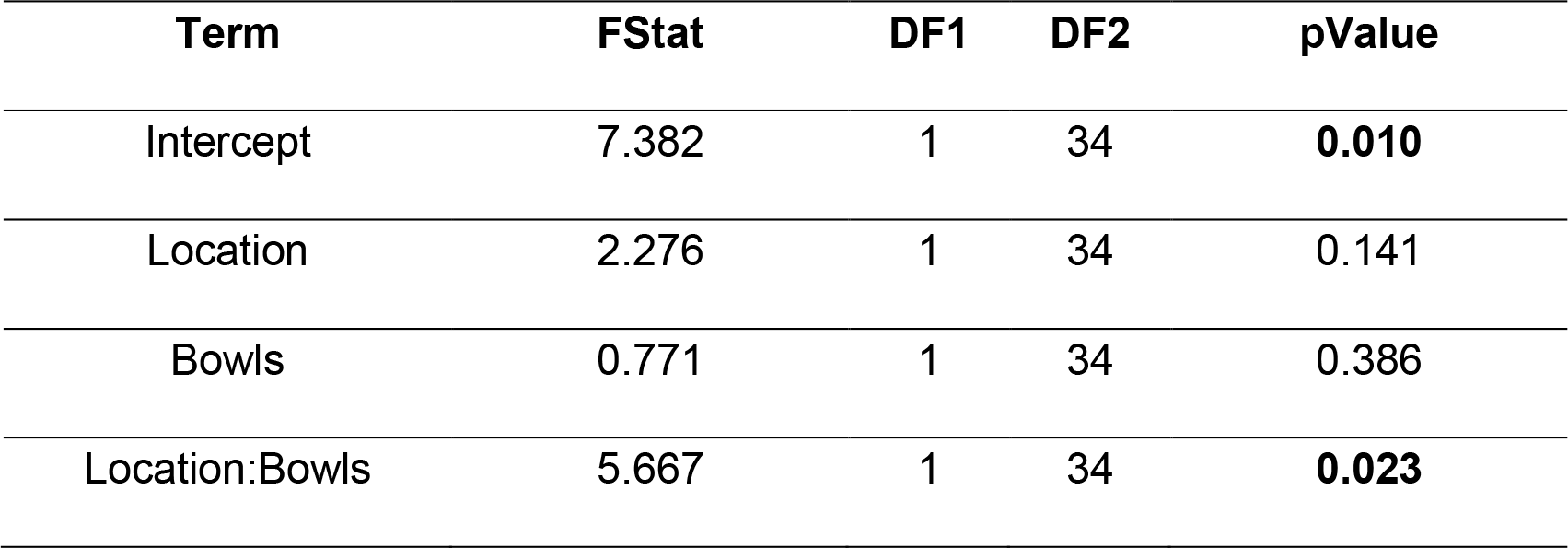
Results from linear mixed-effect model fit. ANOVA table for the fixed effects in the model K ∼ 1 + Location + Bowls + Location:Bowls + (1|Group).

**Figure S1.**
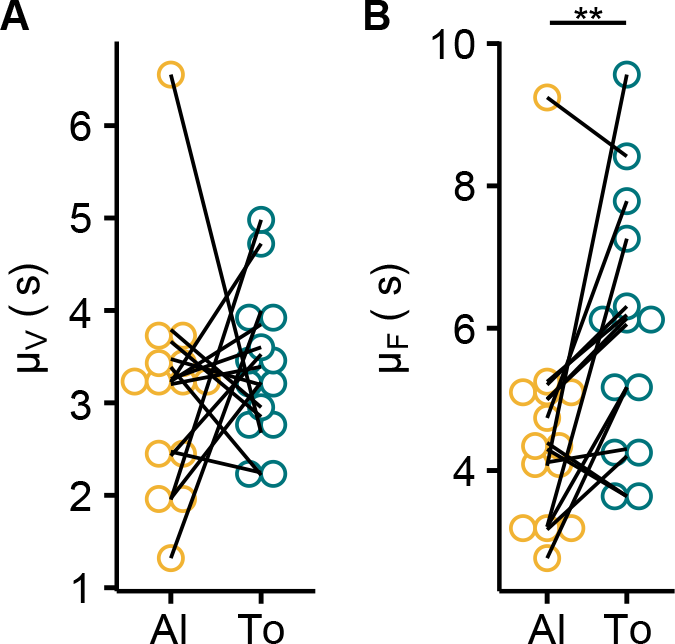
Contributions of vigilant and feeding durations to the increase in time period seen when marmosets are together. Plots compare the fit parameters of vigilance **(A)** and feeding durations **(B)** in the alone (Al) and together (To) conditions. Each point is an individual (n=14 individuals). Individuals belonging to the same group are connected by lines. **p<0.01, two-sided Wilcoxon signed-rank test.

**Figure S2.**
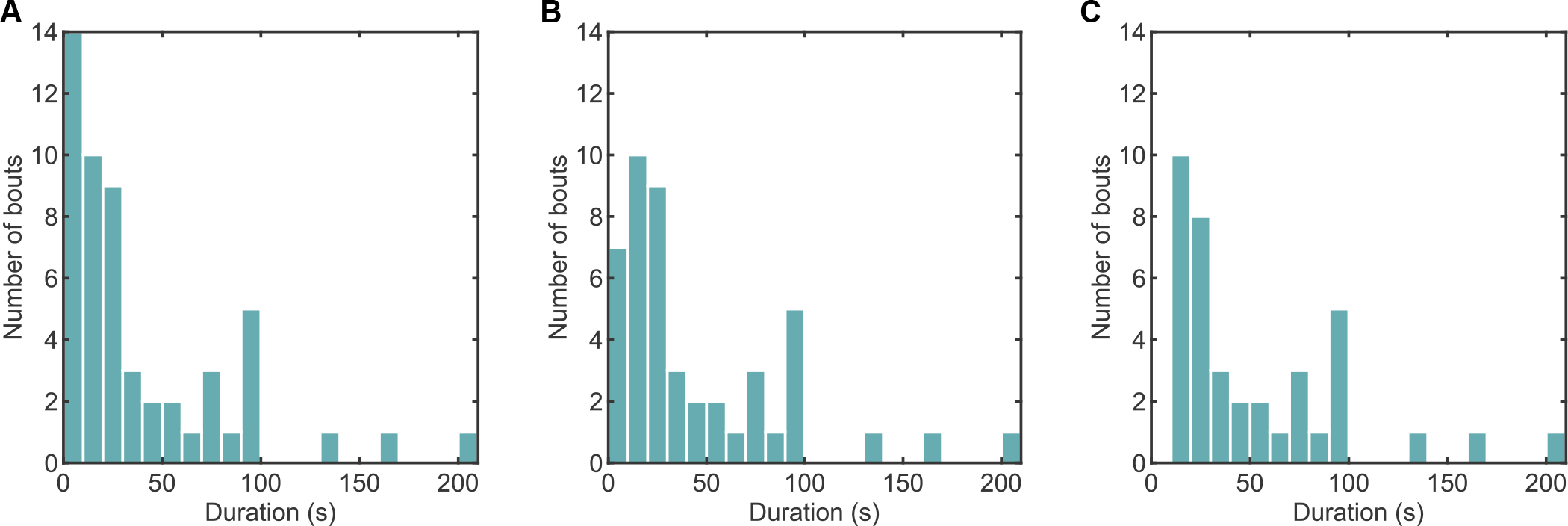
Distributions of bout durations. Histograms for all bouts in the data **(A**), bouts for which the Kuramoto model could be fit **(B)**, and analysed bouts **(C)**.

**Figure S3.**
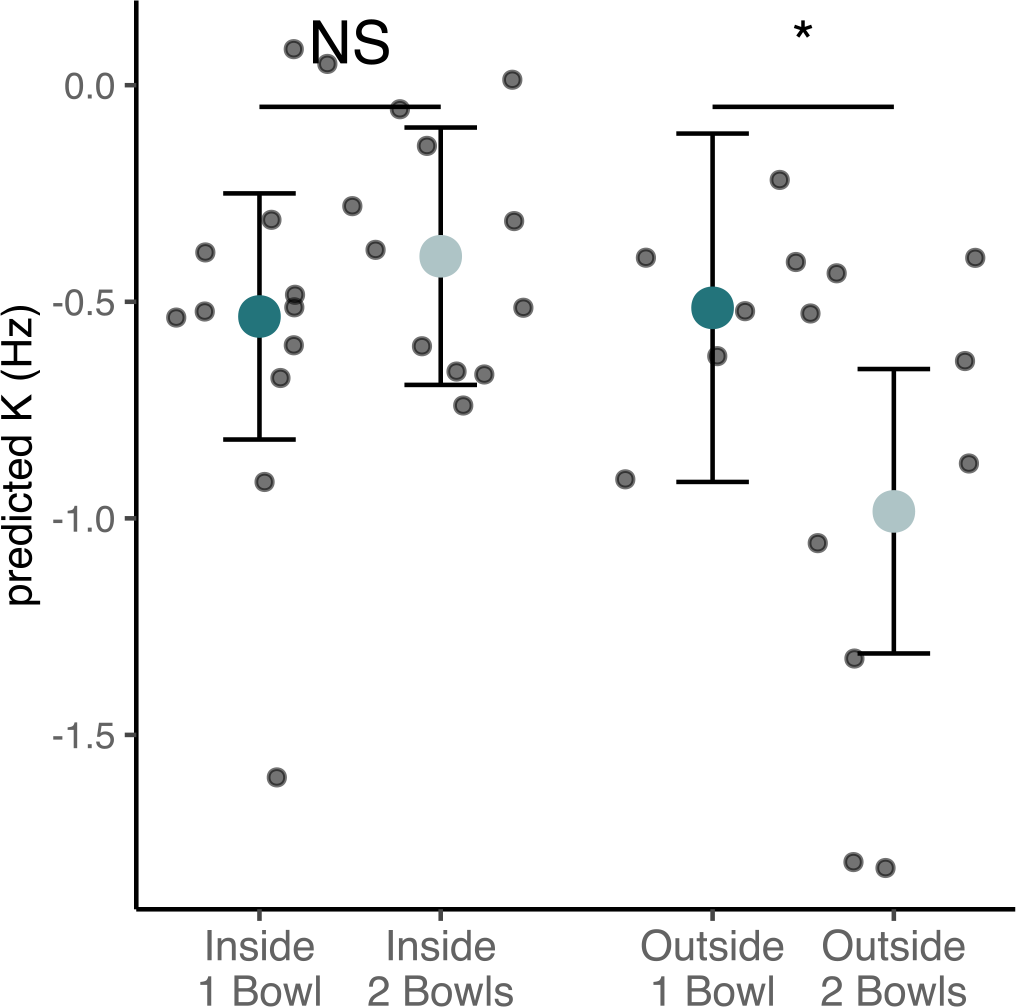
Effect of interaction between location and number of bowls on the coupling constant (K). Each point is a behavioural bout (n=38 bouts). Dots with error bars depict the estimated means and 95% confidence intervals. *p<0.05, NS=p>0.05, post-hoc comparisons of estimated means (Welch’s t-test).

## References

1. Greenfield MD, Merker B. 2023 Coordinated rhythms in animal species, including humans: entrainment from bushcricket chorusing to the philharmonic orchestra. Neuroscience & Biobehavioral Reviews 153, 105382. (doi:10.1016/j.neubiorev.2023.105382)

2. Verga L, Kotz SA, Ravignani A. 2023 The evolution of social timing. Physics of Life Reviews 46, 131–151. (doi:10.1016/j.plrev.2023.06.006)

3. Henry MJ, Cook PF, de Reus K, Nityananda V, Rouse AA, Kotz SA. 2021 An ecological approach to measuring synchronization abilities across the animal kingdom. Philosophical Transactions of the Royal Society B: Biological Sciences 376, 20200336. (doi:10.1098/rstb.2020.0336)

4. Buck J. 1988 Synchronous rhythmic flashing of fireflies. II. The Quarterly review of biology 63, 265–289.

5. Buck J, Buck E. 1968 Mechanism of rhythmic synchronous flashing of fireflies. Science 159, 1319–1327. (doi:10.1126/science.159.3821.1319)

6. Grafe TU. 1996 The function of call alternation in the african reed frog (Hyperolius marmoratus): precise call timing prevents auditory masking. Behav Ecol Sociobiol 38, 149–158. (doi:10.1007/s002650050227)

7. Jones DL, Jones RL, Ratnam R. 2014 Calling dynamics and call synchronization in a local group of unison bout callers. J Comp Physiol A 200, 93–107. (doi:10.1007/s00359-013-0867-x)

8. Zhang W, Yartsev MM. 2019 Correlated Neural Activity across the Brains of Socially Interacting Bats. Cell 178, 413-428.e22. (doi:10.1016/j.cell.2019.05.023)

9. Konvalinka I, Xygalatas D, Bulbulia J, Schjødt U, Jegindø E-M, Wallot S, Van Orden G, Roepstorff A. 2011 Synchronized arousal between performers and related spectators in a fire-walking ritual. Proceedings of the National Academy of Sciences 108, 8514–8519. (doi:10.1073/pnas.1016955108)

10. Cavagna A, Queirós SMD, Giardina I, Stefanini F, Viale M. 2013 Diffusion of individual birds in starling flocks. Proceedings of the Royal Society B: Biological Sciences 280, 20122484. (doi:10.1098/rspb.2012.2484)

11. Pika S, Wilkinson R, Kendrick KH, Vernes SC. 2018 Taking turns: bridging the gap between human and animal communication. Proceedings of the Royal Society B: Biological Sciences 285, 20180598. (doi:10.1098/rspb.2018.0598)

12. Takahashi DY, Narayanan DZ, Ghazanfar AA. 2013 Coupled oscillator dynamics of vocal turntaking in monkeys. Current Biology 23, 2162–2168. (doi:10.1016/j.cub.2013.09.005)

13. Takahashi DY, Fenley AR, Ghazanfar AA. 2016 Early development of turn-taking with parents shapes vocal acoustics in infant marmoset monkeys. Philosophical Transactions of the Royal Society B: Biological Sciences 371, 20150370. (doi:10.1098/rstb.2015.0370)

14. Demartsev V, Strandburg-Peshkin A, Ruffner M, Manser M. 2018 Vocal Turn-Taking in Meerkat Group Calling Sessions. Current Biology 28, 3661-3666.e3. (doi:10.1016/j.cub.2018.09.065)

15. O’Connell-Rodwell CE, Wood JD, Wyman M, Redfield S, Puria S, Hart LA. 2012 Antiphonal vocal bouts associated with departures in free-ranging African elephant family groups (Loxodonta africana). Bioacoustics 21, 215–224. (doi:10.1080/09524622.2012.686166)

16. Fortune ES, Rodríguez C, Li D, Ball GF, Coleman MJ. 2011 Neural Mechanisms for the Coordination of Duet Singing in Wrens. Science 334, 666–670. (doi:10.1126/science.1209867)

17. Fröhlich M, Kuchenbuch P, Müller G, Fruth B, Furuichi T, Wittig RM, Pika S. 2016 Unpeeling the layers of language: Bonobos and chimpanzees engage in cooperative turn-taking sequences. Sci Rep 6, 25887. (doi:10.1038/srep25887)

18. Heggli OA, Cabral J, Konvalinka I, Vuust P, Kringelbach ML. 2019 A Kuramoto model of self-other integration across interpersonal synchronization strategies. PLOS Computational Biology 15, e1007422. (doi:10.1371/journal.pcbi.1007422)

19. Greenfield MD. 1994 Synchronous and alternating choruses in insects and anurans: common mechanisms and diverse functions. American Zoologist 34, 605–615. (doi:10.1093/icb/34.6.605)

20. Jones MDR. 1966 The Acoustic Behaviour of the Bush Cricket Pholidoptera Griseoaptera: I. Alternation, Synchronism and Rivalry Between Males. Journal of Experimental Biology 45, 15–30. (doi:10.1242/jeb.45.1.15)

21. Fulton BB. 1934 Rhythm, Synchronism, and Alternation in the Stridulation of Orthoptera. Journal of the Elisha Mitchell Scientific Society 50, 263–267.

22. Alexander RMcN, Langman VA, Jayes AS. 1977 Fast locomotion of some African ungulates. Journal of Zoology 183, 291–300. (doi:10.1111/j.1469-7998.1977.tb04188.x)

23. Gatesy SM, Biewener AA. 1991 Bipedal locomotion: effects of speed, size and limb posture in birds and humans. Journal of Zoology 224, 127–147. (doi:10.1111/j.1469-7998.1991.tb04794.x)

24. Bouwer FL, Nityananda V, Rouse AA, ten Cate C. 2021 Rhythmic abilities in humans and nonhuman animals: a review and recommendations from a methodological perspective. Philosophical Transactions of the Royal Society B: Biological Sciences 376, 20200335. (doi:10.1098/rstb.2020.0335)

25. Celma-Miralles A, Toro JM. 2020 Discrimination of temporal regularity in rats (rattus norvegicus) and humans (homo sapiens). Journal of Comparative Psychology 134, 3–10. (doi:10.1037/com0000202)

26. Humpal J, Cynx JA. 1984 Discrimination of temporal components of acoustic patterns by birds. Annals of the New York Academy of Sciences 423, 600–602. (doi:10.1111/j.1749-6632.1984.tb23466.x)

27. Hulse SH, Humpal J, Cynx J. 1984 Discrimination and generalization of rhythmic and arrhythmic sound patterns by european starlings (Sturnus vulgaris). Music Perception 1.

28. van der Aa J, Honing H, ten Cate C. 2015 The perception of regularity in an isochronous stimulus in zebra finches (Taeniopygia guttata) and humans. Behavioural Processes 115, 37–45. (doi:10.1016/j.beproc.2015.02.018)

29. Hagmann CE, Cook RG. 2010 Testing meter, rhythm, and tempo discriminations in pigeons. Behavioural Processes 85, 99–110. (doi:10.1016/j.beproc.2010.06.015)

30. Gerhardt HC. 1991 Female mate choice in treefrogs: static and dynamic acoustic criteria. Animal Behaviour 42, 615–635. (doi:10.1016/S0003-3472(05)80245-3)

31. Doherty JA. 1985 Phonotaxis in the cricket, Gryllus bimaculatus DeGeer: comparisons of choice and no-choice paradigms. J. Comp. Physiol. 157, 279–289. (doi:10.1007/BF00618118)

32. Zarco W, Merchant H, Prado L, Mendez JC. 2009 Subsecond timing in primates: comparison of interval production between human subjects and rhesus monkeys. Journal of Neurophysiology 102, 3191–3202. (doi:10.1152/jn.00066.2009)

33. Hasegawa A, Okanoya K, Hasegawa T, Seki Y. 2011 Rhythmic synchronization tapping to an audio–visual metronome in budgerigars. Sci Rep 1, 120. (doi:10.1038/srep00120)

34. Gámez J, Yc K, Ayala YA, Dotov D, Prado L, Merchant H. 2018 Predictive rhythmic tapping to isochronous and tempo changing metronomes in the nonhuman primate. Annals of the New York Academy of Sciences 1423, 396–414. (doi:10.1111/nyas.13671)

35. Takeya R, Patel AD, Tanaka M. 2018 Temporal generalization of synchronized saccades beyond the trained range in monkeys. Frontiers in Psychology 9.

36. Katsu N, Yuki S, Okanoya K. 2021 Production of regular rhythm induced by external stimuli in rats. Anim Cogn 24, 1133–1141. (doi:10.1007/s10071-021-01505-4)

37. Seki Y, Tomyta K. 2019 Effects of metronomic sounds on a self-paced tapping task in budgerigars and humans. Current Zoology 65, 121–128. (doi:10.1093/cz/zoy075)

38. Rouse AA, Cook PF, Large EW, Reichmuth C. 2016 Beat keeping in a sea lion as coupled oscillation: implications for comparative understanding of human rhythm. Frontiers in Neuroscience 10.

39. Cook P, Rouse A, Wilson M, Reichmuth C. 2013 A california sea lion (Zalophus californianus) can keep the beat: motor entrainment to rhythmic auditory stimuli in a non vocal mimic. Journal of Comparative Psychology 127, 412–427. (doi:10.1037/a0032345)

40. Patel AD, Iversen JR, Bregman MR, Schulz I. 2009 Experimental evidence for synchronization to a musical beat in a nonhuman animal. Current Biology 19, 827–830. (doi:10.1016/j.cub.2009.03.038)

41. Schachner A, Brady TF, Pepperberg IM, Hauser MD. 2009 Spontaneous motor entrainment to music in multiple vocal mimicking species. Current Biology 19, 831–836. (doi:10.1016/j.cub.2009.03.061)

42. Schweinfurth MK, Baldridge DB, Finnerty K, Call J, Knoblich GK. 2022 Inter-individual coordination in walking chimpanzees. Current Biology 32, 5138-5143.e3. (doi:10.1016/j.cub.2022.09.059)

43. Pulliam HR. 1973 On the advantages of flocking. Journal of Theoretical Biology 38.

44. Scannell J, Roberts G, Lazarus J. 2001 Prey scan at random to evade observant predators. Proc. R. Soc. Lond. B 268, 541–547. (doi:10.1098/rspb.2000.1388)

45. Allan ATL, Hill RA. 2018 What have we been looking at? A call for consistency in studies of primate vigilance. Am J Phys Anthropol 165, 4–22. (doi:10.1002/ajpa.23381)

46. Beauchamp G. 2017 Difficulties in monitoring conspecifics mediate the effects of visual obstruction on the level and synchronization of vigilance. Front. Ecol. Evol. 5. (doi:10.3389/fevo.2017.00012)

47. Novčić, Mlakar MM, idović Z, Hauber ME. 2023 Black-headed gulls synchronize vigilance with their nearest neighbor irrespective of the neighbor’s relative position. Ethology 129, 146–155. (doi:10.1111/eth.13353)

48. Pays O, Sirot E, Fritz H. 2012 Collective vigilance in the greater kudu: towards a better understanding of synchronization patterns: vigilance in gregarious prey species. Ethology 118, 1–9. (doi:10.1111/j.1439-0310.2011.01974.x)

49. Pays O, Goulard M, Blomberg SP, Goldizen AW, Sirot E, Jarman PJ. 2009 The effect of social facilitation on vigilance in the eastern gray kangaroo, Macropus giganteus. Behavioral Ecology 20, 469–477. (doi:10.1093/beheco/arp019)

50. Öst M, Tierala T. 2011 Synchronized vigilance while feeding in common eider brood-rearing coalitions. Behavioral Ecology 22, 378–384.

51. Rodríguez–Gironés MA, Vásquez RA. 2002 Evolutionary stability of vigilance coordination among social foragers. Proc. R. Soc. Lond. B 269, 1803–1810. (doi:10.1098/rspb.2002.2043)

52. Bednekoff PA. 2015 Sentinel behavior: a review and prospectus. In Advances in the Study of Behavior, Academic Press.

53. Manser MB. 1999 Response of foraging group members to sentinel calls in suricates, Suricata suricatta. Proceedings of the Royal Society of London. Series B: Biological Sciences 266, 1013–1019. (doi:10.1098/rspb.1999.0737)

54. Ostreiher R, Mundry R, Heifetz A. 2021 On the self-regulation of sentinel activity among Arabian babbler groupmates. Animal Behaviour 173, 81–92. (doi:10.1016/j.anbehav.2021.01.002)

55. Ferrari SF. 2009 Predation risk and antipredator strategies. In South American Primates: Comparative Perspectives in the Study of Behavior, Ecology, and Conservation (eds PA Garber, A Estrada, JC Bicca-Marques, EW Heymann, KB Strier), pp. 251–277. New York, NY: Springer. (doi:10.1007/978-0-387-78705-3_10)

56. Stafford BJ, Ferreira FM. 1995 Predation attempts on callitrichids in the Atlantic coastal rain forest of Brazil. Folia primatologica 65, 229–233.

57. Teixeira DS, dos Santos E, Leal SG, de Jesus AK, Vargas WP, Dutra I, Barros M. 2016 Fatal attack on black-tufted-ear marmosets (Callithrix penicillata) by a Boa constrictor: a simultaneous assault on two juvenile monkeys. Primates 57, 123–127. (doi:10.1007/s10329-015-0495-x)

58. Vasquez MRO, Heymann EW. 2001 Crested eagle (Morphnus guianensis) predation on infant tamarins (Saguinus mystax and Saguinus fuscicollis, Callitrichinae). FPR 72, 301–303. (doi:10.1159/000049952)

59. Caine NG. 1998 Cutting costs in response to predatory threat by Geoffroy’s marmosets (Callithrix geoffroyi). American Journal of Primatology 46, 187–196.

60. Gosselin-Ildari AD, Koenig A. 2012 The effects of group size and reproductive status on vigilance in captive Callithrix jacchus. American Journal of Primatology 74, 613–621. (doi:10.1002/aj22013)

61. Koenig A. 1994 Random scan, sentinels or sentinel system? A study in captive common marmosets (Callithrix jacchus). In Social development, learning and behaviour, pp. 69–76. Strasbourg: Université Louis Pasteur.

62. Koenig A. 1998 Visual scanning by common marmosets (Callithrix jacchus): functional aspects and the special role of adult males. Primates 39, 85–90. (doi:10.1007/BF02557746)

63. Brügger RK, Willems EP, Burkart JM. 2023 Looking out for each other: coordination and turn taking in common marmoset vigilance. Animal Behaviour 196, 183–199. (doi:10.1016/j.anbehav.2022.11.007)

64. Kuramoto Y, Kuramoto Y. 1984 Chemical turbulence. Springer.

65. Kuramoto Y. 1975 Self-entrainment of a population of coupled non-linear oscillators. In International Symposium on Mathematical Problems in Theoretical Physics: January 23–29, 1975, Kyoto University, Kyoto/Japan, pp. 420–422. Springer.

66. Strogatz SH. 1997 Spontaneous synchronization in nature. In Proceedings of International Frequency Control Symposium, pp. 2–4. (doi:10.1109/FREQ.1997.638513)

67. Sarfati R, Joshi K, Martin O, Hayes JC, Iyer-Biswas S, Peleg O. 2023 Emergent periodicity in the collective synchronous flashing of fireflies. Elife 12, e78908.

68. Berkowitz A. 2019 Expanding our horizons: central pattern generation in the context of complex activity sequences. Journal of Experimental Biology 222, jeb192054. (doi:10.1242/jeb.192054)

69. Ruch H, Zürcher Y, Burkart JM. 2018 The function and mechanism of vocal accommodation in humans and other primates. Biological Reviews 93, 996–1013.

70. Zürcher Y, Willems EP, Burkart JM. 2019 Are dialects socially learned in marmoset monkeys? Evidence from translocation experiments. PLOS ONE 14, e0222486. (doi:10.1371/journal.pone.0222486)

71. Phaniraj N, Wierucka K, Burkart JM. 2023 Dynamic vocal learning in adult marmoset monkeys., 2023.09.22.559020. (doi:10.1101/2023.09.22.559020)

72. Zürcher Y, Willems EP, Burkart JM. 2021 Trade-offs between vocal accommodation and individual recognisability in common marmoset vocalizations. Scientific Reports 11, 1–10.

73. Boinski S, Kauffman L, Westoll A, Stickler CM, Cropp S, Ehmke E. 2003 Are vigilance, risk from avian predators and group size consequences of habitat structure? A comparison of three species of squirrel monkey (Saimiri oerstedii, S. boliviensis, and S. sciureus). Behaviour 140, 1421–1467.

74. Strogatz SH. 2018 Nonlinear dynamics and chaos with student solutions manual: With applications to physics, biology, chemistry, and engineering. CRC press.

75. Greenfield MD, Roizen I. 1993 Katydid synchronous chorusing is an evolutionarily stable outcome of female choice. Nature 364, 618–620. (doi:10.1038/364618a0)

76. Miss FM, Burkart JM. 2018 Corepresentation during joint action in marmoset monkeys (Callithrix jacchus). Psychol Sci 29, 984–995. (doi:10.1177/0956797618772046)

77. Cronin KA, Kurian AV, Snowdon CT. 2005 Cooperative problem solving in a cooperatively breeding primate (Saguinus oedipus). Animal Behaviour 69, 133–142. (doi:10.1016/j.anbehav.2004.02.024)

78. Miller CT, Mandel K, Wang X. 2010 The communicative content of the common marmoset phee call during antiphonal calling. American Journal of Primatology 72, 974–980. (doi:10.1002/ajp.20854)

79. Miller CT, Wang X. 2006 Sensory-motor interactions modulate a primate vocal behavior: antiphonal calling in common marmosets. J Comp Physiol A 192, 27–38. (doi:10.1007/s00359-005-0043-z)

80. Finkenwirth C, van Schaik C, Ziegler TE, Burkart JM. 2015 Strongly bonded family members in common marmosets show synchronized fluctuations in oxytocin. Physiology & Behavior 151, 246–251. (doi:10.1016/j.physbeh.2015.07.034)

81. Cronin KA, Kurian AV, Snowdon CT. 2005 Cooperative problem solving in a cooperatively breeding primate (Saguinus oedipus). Animal Behaviour 69, 133–142.

82. Faust LF. 2010 Natural history and flash repertoire of the synchronous firefly Photinus carolinus (Coleoptera: Lampyridae) in the Great Smoky Mountains National Park. Florida Entomologist, 208–217.

