## Supplementary materials for "Marmosets mutually compensate for differences in rhythms when coordinating vigilance"

**Table S1. Individuals included in study.** List of all individuals involved in the study detailing their age, status and time as well as order of testing. i = inside; o = outside; 1 = one food bowl; 2 = two food bowls

| **Data collection period** | 03–31 May 2018 | | | | |
|---|---|---|---|---|---|
| **Group** | **Individual** | **Sex** | **Status** | **Birth date** | **Conditions (in order of testing)** |
| Garetta | Garetta | f | breeder | 11 June 2009 | i2; o2; o1; i1 |
| | Nuno | m | breeder | 23 May 2013 | |
| Gaviota | Gaviota | f | breeder | 03 Oct. 2006 | o1; o2; i2; i1 |
| | Kapi | m | breeder | 01 Sept. 2002 | |
| Grappa | Grappa | f | breeder | 17 Mar. 2015 | o1; o2; i1; i2 |
| | Craken | m | breeder | 10 Oct. 2013 | |
| | Ginger | f | infant | 04 May 2018 | *(no vigilance coded)* |
| | Guapa | f | infant | 04 May 2018 | *(no vigilance coded)* |
| Greta | Greta | f | breeder | 11 Oct. 2011 | i1; o1; o2; i2 |
| | Mars | m | breeder | 11 July 2012 | |
| Lilly | Lilly | f | breeder | 03 July 2012 | i1; o1; i2; o2 |
| | Nando | m | breeder | 23 May 2013 | |
| Vesta | Vesta | f | helper | 05 Oct. 2004 | o2; o1; i2; i1 |
| | Vito | m | helper | 30 May 2006 | |
| Wisconsin | Wisconsin | f | breeder | 30 Aug. 2013 | o2; i1; i2; o2 |
| | Tabor | m | breeder | 30 Oct. 2008 | |

**Table S2. Ethogram with definitions for all behaviours/categories coded from video material.** Table taken from Brügger et al. 2023.

| **Behaviour** | **Definition** |
|---|---|
| Vigilance | All looking behaviour (≥ 1s) directed over arm's reach and *not* toward a group member or the substrate the animal is sitting on (following Allan & Hill, 2018) |
| Out of sight | All looking behaviour (≥ 1s) where the line of sight is not clearly discernible, but the animal could potentially be vigilant |
| Feeding | Since the mash can only be eaten when licking it out of the bowl, the feeding starts with the first frame of the subject's head inside the cup, meaning the upper corner of the white ear tufts is no longer visible and it ends with the first frame of the edge of the ear tufts becoming visible at the upper edge of the food bowl |
| **Location** | **Definition** |
| Inside /outside | *Location "inside":*<br>All animals can a priori only be inside: not outside.<br>*Location "outside":*<br>Animals access the outdoor enclosure via a tube from the inside enclosure. They have access to part of the tube where they cannot see the outside enclosure and are thus considered "inside" until the first frame where all limbs are fully outside of the tube and as soon as one limb is inside the tube and "outside" in all other situations. |
| On basket | Animals are considered on feeding basket at the front of the enclosure (inside/outside) starting with the first frame where one limb is on the feeding basket until the last frame where a limb is visible on feeding basket. |

**Table S3. Results from linear mixed-effect model fit.** ANOVA table for the fixed effects in the model K ~ 1 + Location + Bowls + Location:Bowls + (1|Group).

| Term | FStat | DF1 | DF2 | pValue |
|---|---|---|---|---|
| Intercept | 7.382 | 1 | 34 | **0.010** |
| Location | 2.276 | 1 | 34 | 0.141 |
| Bowls | 0.771 | 1 | 34 | 0.386 |
| Location:Bowls | 5.667 | 1 | 34 | **0.023** |

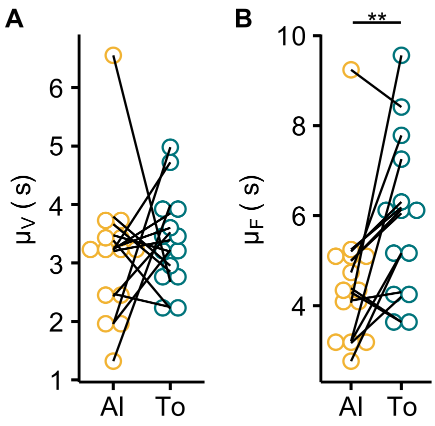


**Figure S1. Contributions of vigilant and feeding durations to the increase in time period seen when marmosets are together.** Plots compare the fit parameters of vigilance **(A)** and feeding durations **(B)** in the alone (Al) and together (To) conditions. Each point is an individual (n=14 individuals). Individuals belonging to the same group are connected by lines. **p<0.01, two-sided Wilcoxon signed-rank test.

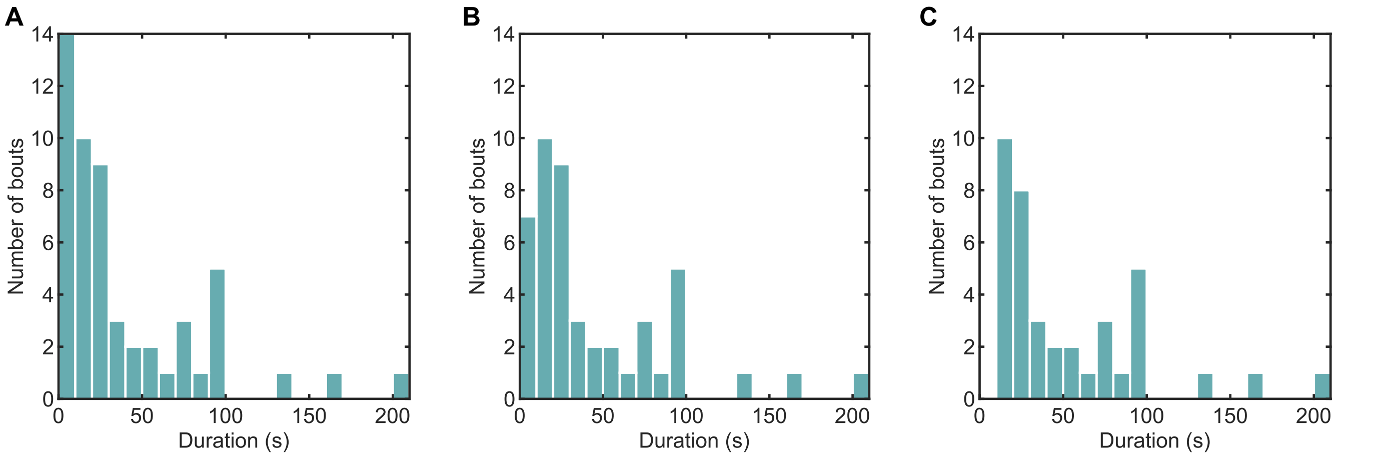


**Figure S2. Distributions of bout durations.** Histograms for all bouts in the data **(A)**, bouts for which the Kuramoto model could be fit **(B)**, and analysed bouts **(C)**.

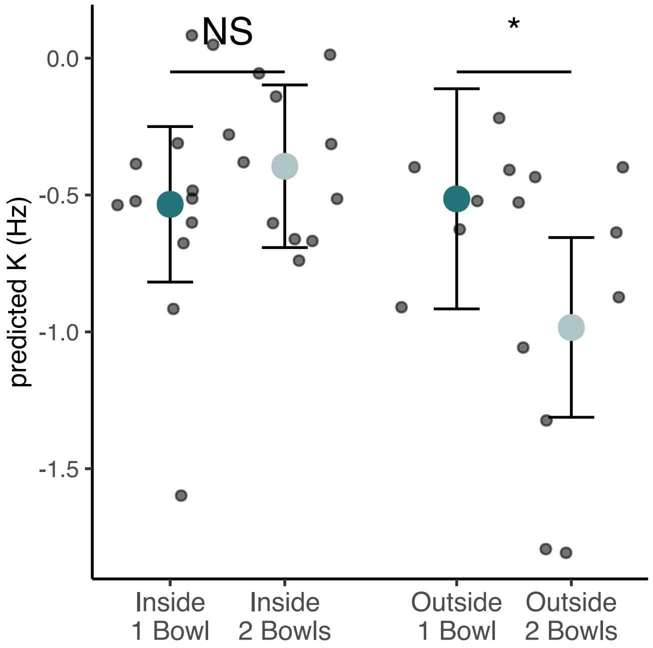


**Figure S3. Effect of interaction between location and number of bowls on the coupling constant (K)**. Each point is a behavioural bout (n=38 bouts). Dots with error bars depict the estimated means and 95% confidence intervals. *p<0.05, NS=p>0.05, post-hoc comparisons of estimated means (Welch's t-test).
